# Antigen reactivity defines tissue-resident memory and exhausted T cells in tumours

**DOI:** 10.1101/2025.07.23.666465

**Authors:** Thomas N. Burn, Jan Schröder, Luke C. Gandolfo, Maleika Osman, Elanor N. Wainwright, Enid Y. N. Lam, Keely M. McDonald, Rachel B. Evans, Shihan Li, Daniel Rawlinson, Lachlan Dryburgh, Ali Zaid, Zoltan Maliga, Dominick Schienstock, Philippa Meiser, Hyun Jae Lee, Hongjin Lai, Marcela L. Moreira, Pirooz Zareie, Louis H-Y. Lee, Lutfi Huq, Susan N. Christo, Justine J. W. Seow, Keith A. Ching, Stéphane M Guillaume, Kathy Knezevic, Simone L. Park, Maximilien Evrard, Jason Waithman, Thomas Gebhardt, Scott N. Mueller, Georgina E. Riddiough, Marcos V. Perini, Simon C. H. Tsao, Terence P. Speed, Peter K. Sorger, Sherene Loi, Francis R. Carbone, Stephanie Gras, Timothy S. Fisher, Bas J. Baaten, Mark A. Dawson, Laura K. Mackay

## Abstract

CD8^+^ T cells are a key weapon in the therapeutic armamentarium against cancer. While CD8^+^CD103^+^ T cells with a tissue-resident memory T (T_RM_) cell phenotype have been favourably correlated with patient prognoses^1–6^, the tumour microenvironment also contains dysfunctional exhausted T (T_EX_) cells that exhibit a myriad of T_RM_-like features, leading to conflation of these two populations. Here, we deconvolute T_RM_ and T_EX_ cells within the intratumoural CD8^+^CD103^+^ T cell pool across human cancers, ascribing markers and gene signatures that distinguish these CD8^+^ populations and enable their functional distinction. We found that while T_RM_ cells exhibit superior functionality and are associated with long-term survival post-tumour resection, they are not associated with responsiveness to immune checkpoint blockade. Deconvolution of the two populations showed that tumour-associated T_EX_ and T_RM_ cells are clonally distinct, with the latter comprising both tumour-independent bystanders and tumour-specific cells segregated from their cognate antigen. Intratumoural T_RM_ cells can be forced towards an exhausted fate when chronic antigen stimulation occurs, arguing that the presence or absence of continuous antigen exposure within the microenvironment is the key distinction between respective tumour-associated T_EX_ and T_RM_ populations. These results suggest unique roles for T_RM_ and T_EX_ cells in tumour control, underscoring the need for distinct strategies to harness these T cell populations in novel cancer therapies.

## Main text

T cell-mediated tumour control is a key facet of cancer immunotherapy and pinpointing the most effective T cell subtypes for therapeutic targeting is a critical area of ongoing research. Two subsets of significant interest to the cancer immunotherapy field are tissue-resident memory T (T_RM_) cells and exhausted T (T_EX_) cells. CD8^+^ T_RM_ cells are a non-recirculating memory T cell population that reside long-term in every organ examined^7–9^. Canonical T_RM_ cells develop following the resolution of infection or inflammation, where they can provide rapid, localised immune protection^10,11^. In contrast, CD8^+^ T_EX_ cells form in the context of chronic infection and cancer, driven by persistent antigen recognition and inflammation^12–14^. T cell exhaustion is characterised by the loss of proliferative capacity and function such that restoration of T cell activity forms the basis of successful immune checkpoint blockade (ICB) therapies. Despite this, while T_EX_ cells may be temporarily reinvigorated following ICB, their long-term fate remains largely unaltered^15^. Most recently, T_EX_-phenotype cells have been shown to become resident in tumours, likely sharing some aspects of the T_RM_ transcriptional program to limit recirculation^16^. Because many studies use genetic signatures to identify T cell subtypes, their transcriptional similarities have resulted in T_RM_ and T_EX_ cells being conflated in the literature. While tumour-associated T_RM_ cells may exist as a potential antipode to the dysfunctional T_EX_ population, their identification within tumours and contribution to cancer control has not been adequately addressed.

### T_EX_ cells share the T_RM_ gene signature

Experimental systems such as parabiosis or transplantation have demonstrated the non-circulating behaviour of T_RM_ cells^10,11,17^. Such experiments in mice led to the identification of T_RM_-associated markers including CD69 and CD103 that distinguish T_RM_ cells from circulating memory T cells (T_CIRCM_), with markers partially cross-validated in humans by their expression on donor-derived T cells present within transplanted organs^18–20^. T_RM_ cells defined by CD69 or CD103 expression are transcriptionally distinct from T_CIRCM_ in humans^7,21,22^ and mice^8,23^. At the core of the T_RM_ gene signature exists a transcriptional program designed to halt T cell migration, which includes downregulated expression of key regulators of T cell recirculation such as *KLF2*, *S1PR1*, and *S1PR5*^24,25^. However, it has recently been shown that T_EX_ cells also cease to migrate and become resident in tumours^16^. Thus, we hypothesised that T_EX_ cells utilise a common transcriptional program to T_RM_ cells to inhibit migration, resulting in considerable transcriptional overlap between these populations. Indeed, we revealed this is the case, with the T_RM_ transcriptional profile derived from acute viral infection models (lymphocytic choriomeningitis virus (LCMV) and herpes simplex virus (HSV))^8^ significantly correlated to the T_EX_ transcriptional profile derived from chronic versus acute LCMV infection^26^ (**Fig. 1a)**. Given this overlap between T_RM_ and T_EX_ cells, we reasoned that T_RM_ gene signatures would identify T_EX_ cells in single-cell RNA sequencing (scRNAseq) datasets. To test this, we utilised published data of CD8^+^ T cells from spleens of mice infected with acute or chronic LCMV^27^. Strikingly, we found the T_RM_ gene signature was most enriched in terminal T_EX_ cells during chronic infection (**Fig. 1b-d, Extended Data Fig 1a**). Further, we analysed CD8^+^ T cells from murine breast cancer (BC) and adjacent tissue^16^, finding the T_RM_ gene signature highly enriched in tumour-specific T_EX_ cells isolated from tumours, similar to virus-specific cells in adjacent tissue (**Fig. 1e-f**). Thus, the utilisation of T_RM_ gene signatures to identify intratumoural T_RM_ cells results in aberrant T_EX_ cell identification.

**Figure 1:**
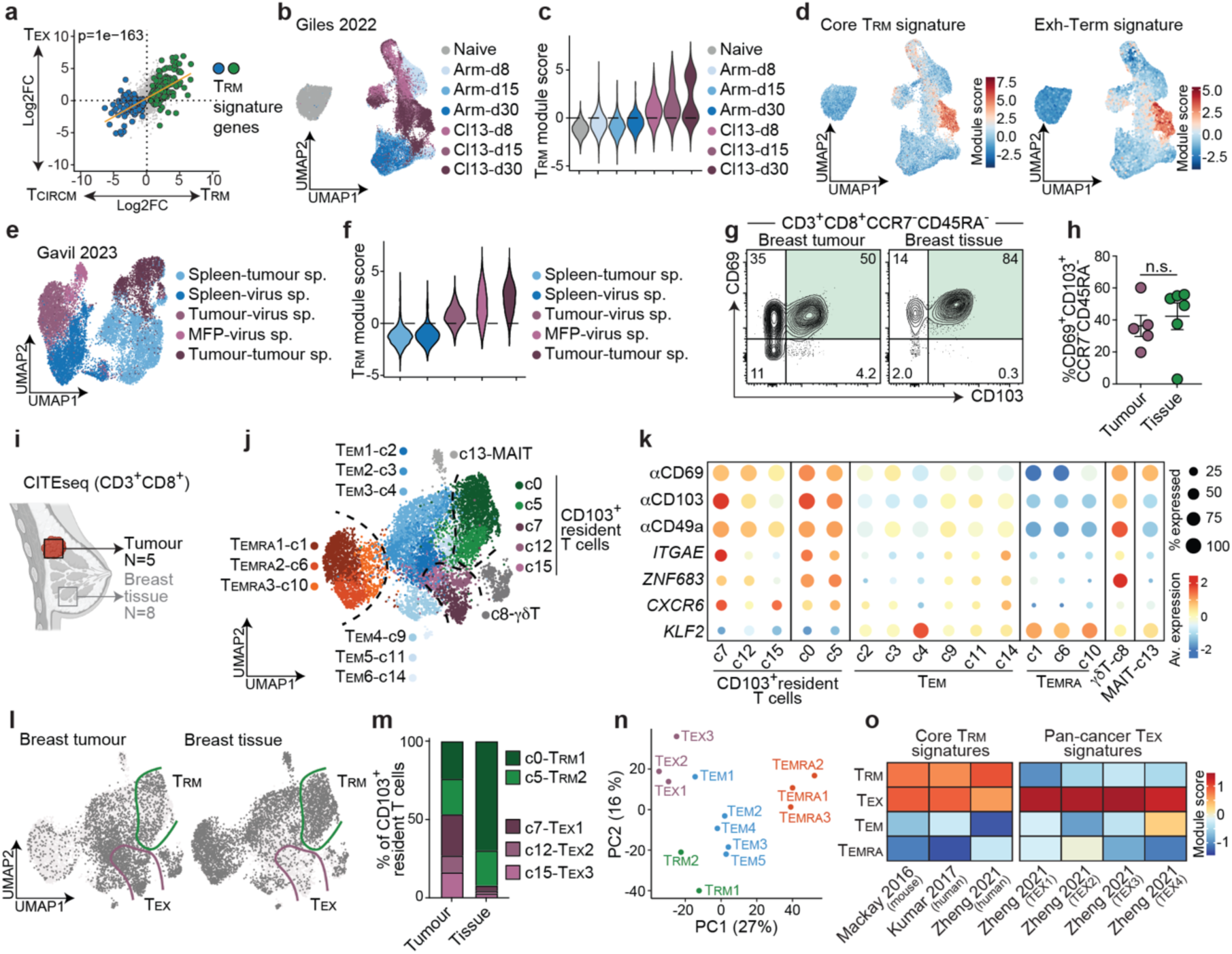
Canonical T_RM_ proteins and gene signatures do not deconvolute T_RM_ and T_EX_ cells. **a**, Log2 fold-change (FC) of differentially expressed T_RM_^8^ and T_EX_^26^ cell genes compared to circulating memory T cells (T_CIRCM_). T_RM_ signature genes^8^ are highlighted (green=up, blue=down). p-value indicates Fisher’s exact test for association. **b-d**, scRNA-seq data from LCMV-specific T cells after acute (Arm) or chronic (Cl13) infection isolated from spleens at d8, d15, or d30 post-infection^27^. **b**, UMAP projection of scRNA-seq data annotated by infection and timepoint, **c,** quantification of T_RM_ module score^8^ and **d**, overlay of core T_RM_^8^and Exh-Term^27^ transcriptional signatures on respective populations. **e-f**, UMAP projection (**e**) and quantification of T_RM_ module score^8^ (**f**) on LCMV-specific P14 (virus-sp) or tumour-specific OT-I T cells from CITEseq analysis of the respective organs following EO771-OVA BC tumours^16^. **g-h**, Flow cytometry of CD8^+^ T cells isolated from BC tumours or breast tissue from N=5 BC patients. **g**, Representative plots and **h**, summary data for %CD69^+^CD103^+^CCR7^-^CD45RA^-^ of CD8^+^ T cells. **i-o**, CITEseq of CD3^+^CD8^+^ T cells from primary BC tumours (N=5) and non-cancerous breast tissue (N=8). **i,** Schematic. **j,** Data was Harmony-integrated, and unified protein and RNA-seq data were represented on weighted nearest neighbours UMAP and coloured by cluster. **k**, Expression of respective cell surface proteins (αCD103, αCD69, αCD49a) and transcripts (*ITGAE, ZNF683, CXCR6, KLF2)* across annotated clusters. **l-m**, CD8^+^ T cells segregated by tissue of origin (**l**), and relative cluster composition of CD103^+^ resident T cells isolated from BC tumours or tissue (**m**). **n**, PCA of pseudobulked clusters annotated in (j). **o,** average module scores of published T_RM_^8,21,28^ and T_EX_^28^ gene signatures by annotated subsets.

Many studies have also identified T_RM_ cells in tumours via CD69 and CD103 co-expression^1–6^. CD69^+^CD103^+^ CD8^+^ T cells are present in both non-cancerous tissue and tumours, the latter of which likely encompasses a mixed population of T_RM_ and T_EX_ cells (**Fig. 1g-h, Extended Data Fig. 1b**). Thus, CD69, CD103, and T_RM_ gene signatures all appear insufficient to distinguish T_RM_ and T_EX_ cells should they coexist. Given this overlap in tissue-residency features, we set out to differentiate T_RM_ and T_EX_ populations in human tumours by formulating two testable assumptions. First, cells enriched for expression of core-residency genes and gene signatures in healthy tissue in the absence of overt infection or inflammation are predominantly bona fide T_RM_ cells, while tumour-derived cells expressing these signatures comprise both T_RM_ and T_EX_ cells. Second, that tumour-derived cells expressing core-residency gene signatures can be further segregated into T_EX_ and T_RM_ cells by the relative presence or absence of ‘exhaustion’ gene-signatures. If there are bona fide T_RM_ cells in tumours, they would be expected to be transcriptionally similar T_RM_ cells to those in associated healthy tissue. To this end, we performed single-cell cellular indexing of transcriptomes and epitopes by sequencing (CITEseq) with TCR profiling on CD3^+^ T cells from human BC tumours and normal breast tissue (**Fig. 1i**). CD8^+^ T cells from BC tumours and breast tissue were distributed over 15 clusters, classified into 5 major T cell subsets (T_EMRA_, T_EM_, MAIT, γ8 T cells, and CD103^+^ resident cells) based on protein and transcriptional profiles (**Fig. 1j-k, Extended Data Fig. 1c**). CD103^+^ “resident T cells” shared expression of canonical residency genes including *ZNF683* (HOBIT) and *CXCR6*, and downregulation of *KLF2*, which controls major tissue egress-promoting gene products^24^ (**Fig. 1k**). As per above, we reasoned that tumour-derived CD103^+^ resident T cells would include both T_RM_ and T_EX_ populations while healthy tissue-derived cells would primarily contain T_RM_ cells. Accordingly, we defined two CD103^+^ clusters (c0, c5) as T_RM_ cells, based on their over-representation (>90%) within healthy tissue, while clusters (c7, c12, c15), primarily found in tumours and largely absent from healthy tissue, were classified as T_EX_ cells (**Fig. 1l-m**). Pseudobulk PCA analysis confirmed the distinction between T_RM_ and T_EX_ clusters, whilst highlighting their similarity in the PC1 axis, driven predominantly by the downregulation of genes associated with T cell egress including *KLF2* (**Fig. 1n, Extended Data Fig. 1d**).

Supporting our annotations, published T_RM_ gene signatures^8,21,28^ were enriched within both T_RM_ and T_EX_ populations while T_EX_ gene signatures^28^ were selectively enriched within T_EX_ cells (**Fig. 1o**). Segregation of T_EX_ and T_RM_ cell populations revealed that while they share expression of CD103, CD69, and *ZNF683* (HOBIT), and the downregulation of *KLF2*, T_EX_ cells expressed higher levels of CD38, CD39, PD-1, CTLA-4, *TIGIT* and *HAVCR2* (TIM3) (**Extended Data Fig. 1e-i**). These data show that T_EX_ cells within human tumours co-opt a residency program, and that commonly utilised T_RM_ cell-associated proteins or gene signatures cannot distinguish between T_RM_ and T_EX_ populations.

### T_RM_ and T_EX_ cells exhibit disparate functional capacities

To disentangle tumour-associated T_RM_ and T_EX_ cells, we developed gene signatures from our BC dataset that could accurately distinguish these populations. Genes were included in the T_RM_ gene signature based on differential expression (DE) in T_RM_ cells compared to other T cell subsets, followed by successive DE analysis between T_RM_ and T_EX_ metaclusters. An analogous approach was used to derive the T_EX_ gene signature (**Fig. 2a-c, Extended Data Fig. 2a-b**). While both T_RM_ and T_EX_ cells were defined by CD103 expression and *KLF2* downregulation **(Fig. 2d, Extended Data Fig. 2c)**, they were further distinguished by expression of markers including CD94, CD161, CD73, CD38, CD101, CD39, *GNLY*, and PD-1 **(Fig. 2e, Extended Data Fig. 2d**). This enabled reliable discrimination of the two populations via cyclic immunofluorescence microscopy (CycIF^29,30^) and flow cytometry, allowing examination of their intratumoural location and functional properties.

**Figure 2:**
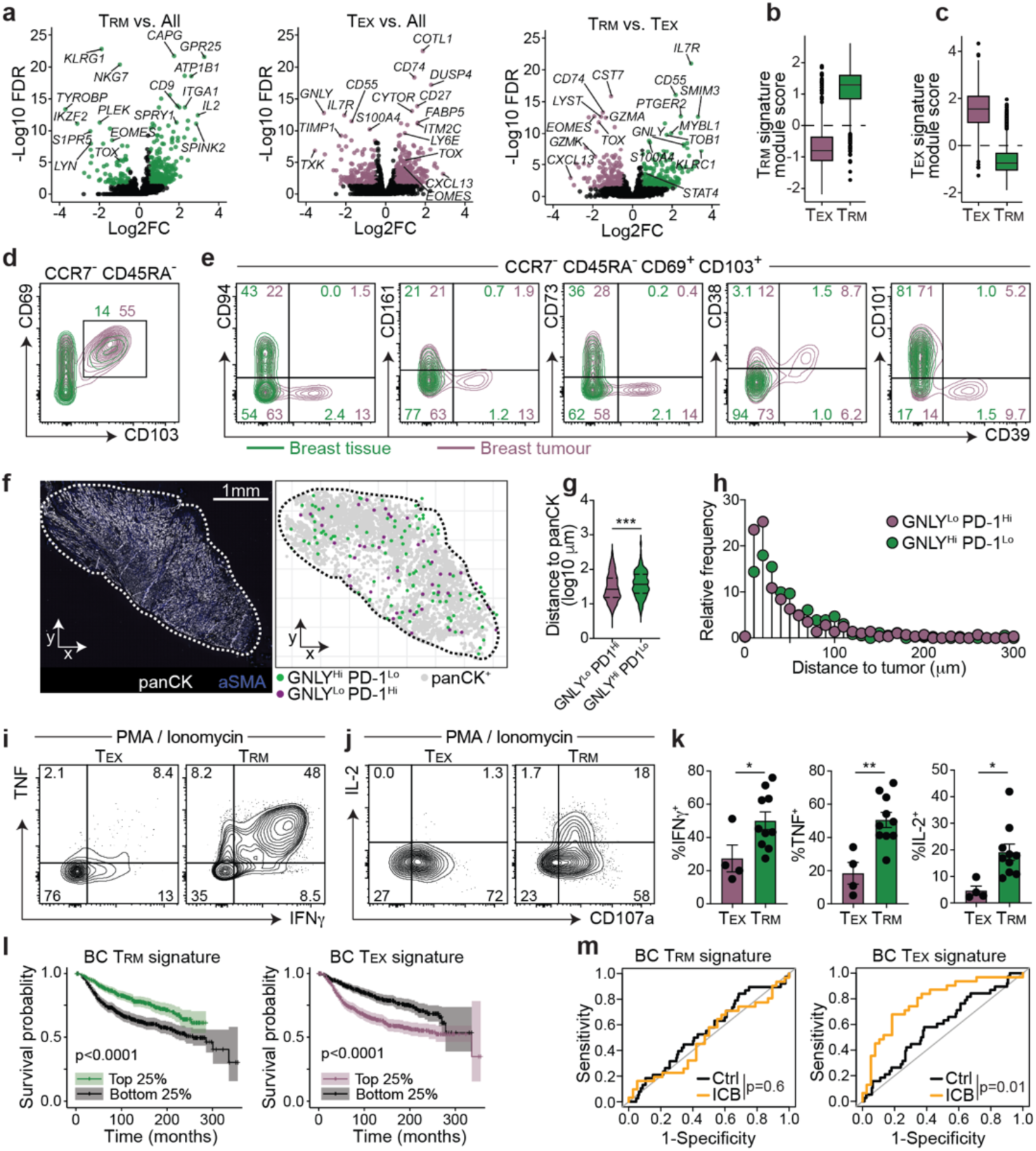
Spatial and functional characterisation of CD103^+^ T_RM_ and T_EX_ cells in tumours. **a,** Volcano plots showing differential expression between T_RM_, T_EX_, and all other subsets in BC dataset. **b-c,** Enrichment of T_RM_ (**b**) and T_EX_ gene signatures (**c**) on labelled subsets. **d-e,** Flow cytometry, concatenated from N=13 donors (N=11 with tumour sample, and N=10 with healthy tissue). **d,** expression of CD69 and CD103 on CCR7^-^CD45RA^-^ CD8^+^ T cells. **e,** expression of T_RM_ and T_EX_-defining proteins on CD69^+^CD103^+^ T cells. **f-h,** CycIF imaging of BC tumours. **f,** representative image of tumour section showing panCK (tumour) and aSMA (stroma) expression, and relative location of respective annotated cell types. Distance to nearest panCK^+^ cell (**g**), and frequency in bins (10μm) segregated by distance of respective cell types to panCK^+^ cells, pooled from N=7 donors (g analysed by t-test, ***p<0.001). Expression of TNF, IFNγ (**i**), IL-2 and CD107a (**j**) on T_EX_ and T_RM_ cells from PMA-ionomycin stimulated CD8^+^ T cells from BC tumours or tissue as per clusters in Extended Data Fig. 3b. **k**. Summary of i-j. Donors contributing a minimum of 10 cells within a cluster (T_RM_ or T_EX_ isolated from either tumour or healthy tissue) were enumerated and plotted (N=4 T_EX_, N=10 T_RM_), analysed by t-test, *p<0.05, **p<0.01. **l,** survival of BC patients from the METABRIC dataset^50^ with the highest (top 25%) T_RM_ or T_EX_ gene signature score enrichments compared to patients with lowest (bottom 25%) gene enrichment scores, plotted on Kaplan-Meier curves with log-rank test. **m,** receiver operating characteristic (ROC) curves from BC patients, either control (Ctrl N=210) or treated with ICB (pembrolizumab/anti-PD-1 N=69) from the iSPY trial^31^. Clinical response to ICB associated with T_RM_ or T_EX_ gene signatures respectively.

Using a 47-marker CycIF panel, we identified CD103^+^ KLF2^-^ CD8^+^ T cells across 7 BC patients **(Extended Data Fig. 2e-f)**. These cells were stratified based on the relative expression of GNLY and PD-1 into GNLY^Hi^ PD-1^Lo^ and GNLY^Lo^ PD-1^Hi^ populations, approximating T_RM_ and T_EX_ cells, respectively **(Extended Data Fig. 2e-g)**. GNLY^Lo^ PD-1^Hi^ cells expressed higher levels of CD39 and LAG3, consistent with an exhausted phenotype **(Extended Data Fig. 2h)**. Unbiased clustering of CD103^+^ KLF2^-^ CD8^+^ T cells reinforced this distinction: GNLY^Hi^ PD-1^Lo^ cells were enriched in cluster c3, which expressed T_RM_-associated markers including CD94, CD7, and NKG2A, while GNLY^Lo^ PD-1^Hi^ cells dominated cluster c1 with increased TIM3, LAG3, and CD39 expression **(Extended Data Fig. 2i-k)**, supporting our *in situ* gating strategy. Spatial analysis revealed that both populations localised near panCK^+^ tumour regions **(Fig. 2f, Extended Data Fig. 2l)**. However, GNLY^Lo^ PD-1^Hi^ (approximating T_EX_) cells were on average significantly closer to panCK^+^ tumour cells, suggesting an increased potential for direct tumour interaction **(Fig. 2g, h)**.

Beyond phenotypic and spatial differences, we also observed distinct functional capacities between T_RM_ and T_EX_ populations. Upon *ex vivo* restimulation, T_RM_ cells exhibited higher production of IFNγ, TNF, and IL-2 and showed elevated expression of granulysin (*GNLY),* while T_EX_ cells expressed more granzyme A (*GZMA*) and granzyme K (*GZMK*) (**Fig. 2i-k**, **Extended Data Fig. 3a-e)**. Moreover, deconvolution of these populations in transcriptomic datasets revealed that enrichment for the T_RM_ gene signature was associated with improved overall survival in BC patients, whereas T_EX_ gene signature enrichment correlated with poorer outcomes (**Fig. 2l**). This association appeared BC-subtype specific, given that triple negative BC (TNBC) survival was best predicted by total CD103^+^ (both T_RM_ and T_EX_) cells, consistent with our prior work (**Extended Data Fig. 3f-h**)^1^. Further, we tested the association of the T_RM_ and T_EX_ signatures with responses to ICB (pembrolizumab/αPD-1) in the I-SPY2 trial^31^, revealing that patients with higher T_EX_ gene signature expression were more likely to achieve a pathologic complete response (pCR) following ICB, whereas elevated T_RM_ signature expression was inconsequential to ICB responsiveness (**Fig. 2m**). These data suggest that while T_RM_ cells are associated with positive prognoses in breast cancer, current ICB therapies exclusively target and enhance T_EX_ cell-mediated anti-tumour responses.

### T_RM_ and T_EX_ gene signatures delineate T_RM_ cells across tumours

To determine the applicability of these BC T_RM_ and T_EX_ gene signatures across tumour types, we next developed signatures from CD8^+^ T cells isolated from liver tumours for comparison. To this end we performed CITEseq on CD8^+^ T cells isolated from liver metastases from colorectal cancer patients, paired non-cancerous liver tissue, and liver tissue from cancer-free donors (**Fig. 3a**). Two CD103^+^ resident (CD69^+^, CD103^+^, *KLF2^-^*) T cell clusters were identified and denoted as T_EX_ and T_RM_ populations as described above, with T_EX_ cells mostly present in tumour-derived tissue and T_RM_ clusters present in both tumour and non-cancerous tissue (**Extended Data Fig. 4a-d**). In line with our findings in BC, we found that both T_RM_ and T_EX_ populations were enriched for published T_RM_ but not T_EX_ gene signatures (**Extended Data Fig. 4e**). Importantly, liver T_RM_ and T_EX_ signatures could accurately identify BC T_RM_ and T_EX_ cells (**Fig. 3b**), and similarly, BC T_RM_ and T_EX_ signatures identified liver T_RM_ and T_EX_ cells, respectively (**Extended Data Fig. 4f**). Overall, BC T_RM_ signature genes were highly enriched in liver T_RM_ cells (and vice versa) which was also true for the respective T_EX_ gene signatures highlighting shared gene expression across tumour-associated T_RM_ and T_EX_ cells from disparate tumour types (**Fig. 3c, Extended Data Fig. 4g**).

**Figure 3:**
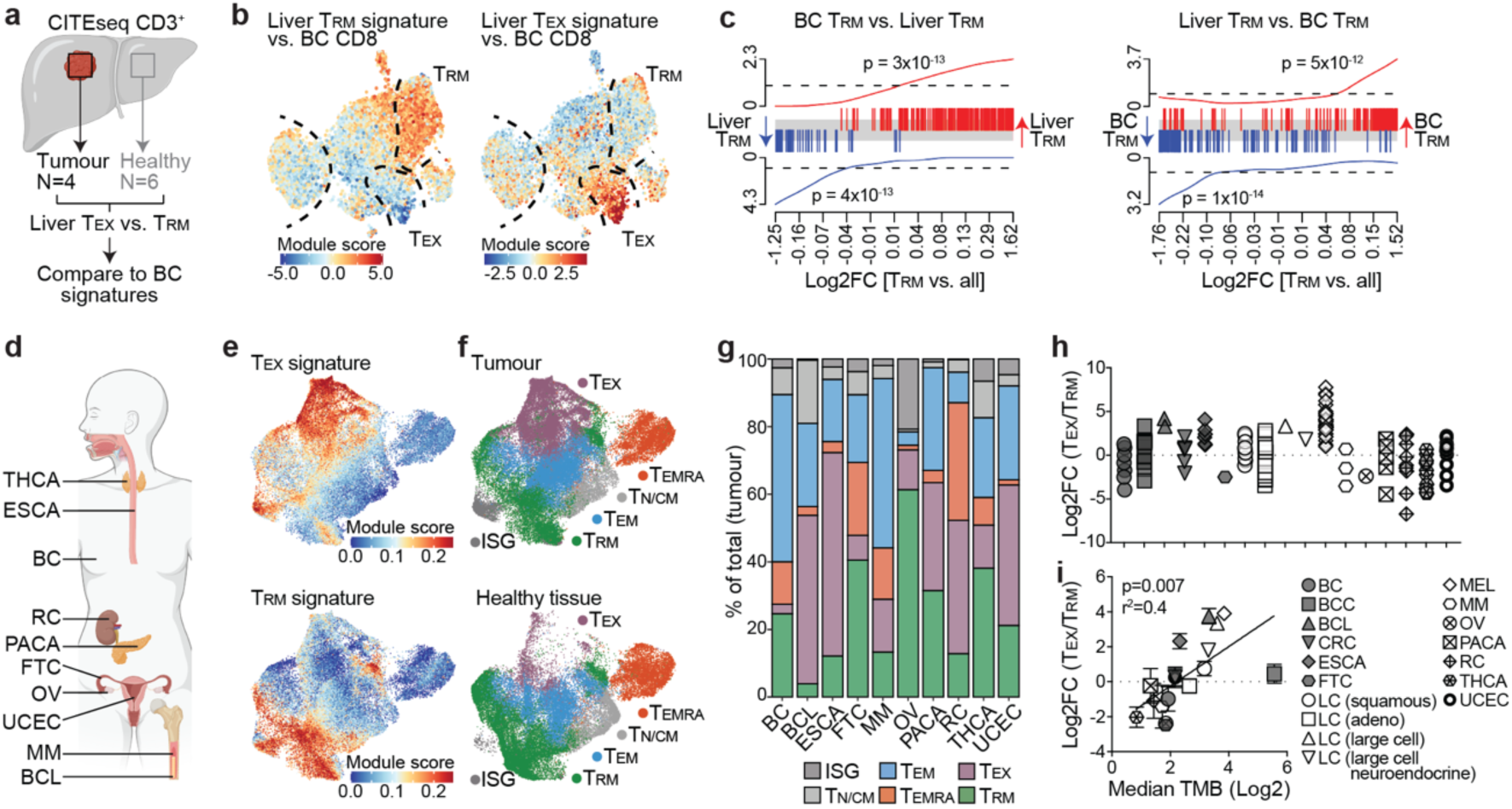
Specific T_RM_ and T_EX_ gene signatures enable T_RM_ and T_EX_ cell identification across tumours. **a,** CITEseq of CD3^+^CD8^+^ T cells isolated from secondary liver tumours (colorectal cancer patients, N=4) and non-cancerous liver tissue (N=6). Liver T_RM_ vs. T_EX_ gene signatures were acquired as described for BC signatures. **b,** Overlay of liver T_RM_ and T_EX_ signature module scores on BC dataset. **c,** Gene-set enrichment analysis of BC T_RM_ vs. liver T_RM_ signatures and vice versa. **d-g,** scRNA-seq of tumour-associated CD8^+^ T cells from various cancers. **d,** Cancers from pan-cancer atlas^28^. **e,** Overlay of refined T_EX_ (top) and T_RM_ (bottom) gene signatures on CD8^+^ T cells on UMAP of pan-cancer scRNA-seq dataset^28^. **f,** Tumour-derived (top) and healthy tissue-derived (bottom) CD8^+^ T cells, and annotated clusters. **g,** Relative frequencies of different CD8^+^ T cell subsets by cancer^28^. **h,** Log2 ratio of T_EX_ to T_RM_ across different cancers^28,40,51–55^. **i,** T_EX_/T_RM_ ratio association with median tumour mutational burden (TMB)^32^. p-value indicates that the slope of the regression line departs from zero. BC (breast), BCC (basal cell carcinoma, BCL (B cell lymphoma), CRC (colorectal cancer), ESCA (esophageal), FTC (fallopian tube), LC (lung cancer), MEL (melanoma), MM (multiple myeloma), OV (ovarian), PACA (pancreatic), RC (renal), THCA (thyroid), UCEC (uterine corpus endometrial).

While the collection of genes within T_RM_ and T_EX_ signatures strongly correlated across BC and liver tumours, the expression of individual genes and surface proteins was not universally consistent. For example, CD94, CD101, CD161 and CD73 were specifically enriched in T_RM_ cells in BC **(Fig. 2e)** but not liver-derived tumours (**Extended Data Fig. 4h**). Therefore, caution must be applied when extrapolating individual genes and proteins across T cell populations in distinct settings. Nonetheless, by focusing on the leading-edge genes, those contributing to the enrichment in each comparison (**Fig. 3c, Extended Data Fig. 4g**), we defined broadly applicable pan-cancer T_RM_ and T_EX_ signatures (**Extended Data Fig. 5a,b**). Using these gene signatures, we could distinguish T_RM_ and T_EX_ cells across a range of tumours from multiple datasets, including a pan-cancer atlas^28^ and additional studies^48–53^ (**Fig. 3d-g, Extended Data Fig. 5c**). A composite of the samples from the pan-cancer atlas showed that T_EX_ cells were predominantly detected in tumour-derived tissue and largely absent from non-cancerous, healthy tissue (**Fig. 3e-f**). When analysed by tumour type, the relative abundance of T_EX_ and T_RM_ cells varied, with melanoma (MEL), oesophageal cancer (ESCA), and B cell lymphoma (BCL) displaying the highest T_EX_ to T_RM_ ratio among the cancers examined (**Fig. 3g-h**). Strikingly, this ratio correlated positively with tumour mutational burden (TMB)^32^, suggesting that neoantigen load may preferentially promote T_EX_ cell formation (**Fig. 3i**). Overall, these data demonstrate our ability to deconvolute T_RM_ and T_EX_ cells across various tumour settings, facilitating a detailed investigation of their functional and developmental differences.

### Tumour-associated T_RM_ are clonally distinct and enriched for virus-specific cells

Given the observed correlation between T_EX_ cell abundance and TMB, we hypothesised that T_EX_ and T_RM_ populations may be clonally distinct, potentially reflecting differences in antigen specificity. To investigate this, we examined clonal overlap among the top 100 expanded TCR clones across T cell subsets in both our BC dataset and the pan-cancer atlas. In both datasets, T_RM_ and T_EX_ cells displayed limited clonal overlap with each other and instead showed greater clonal overlap with T_EM_ cells (**Fig. 4a**). High Jaccard dissimilarity scores supported this, indicating that TCR repertoires of tumour-derived T_RM_, T_EX_ and T_EM_ cells were largely distinct (**Fig. 4b**). Within our BC dataset, expanded clones were occasionally shared across subsets, with most sharing occurring between T_RM_ and T_EM_ or T_EX_ and T_EM_, rather than directly between T_RM_ and T_EX_ **(Fig. 4c)**. Since no public TCRs were present across donors in our BC dataset, liver dataset, or the pan-cancer atlas, we pooled the TCR data for further examination. We assessed clonotype sharing among T_RM_, T_EX_, and T_EM_ cells at various thresholds based on overlapping cell numbers for each clonotype, and found that most clonal overlap among T_RM_, T_EX_, and T_EM_ cells was due to a single shared cell despite significant numbers of expanded clones (**Fig. 4d, Extended Data Fig. 6a**). These data indicate that while a single clone can adopt T_RM_, T_EX_, or T_EM_ phenotypes, there is preferential development of one subset for a given TCR.

**Figure 4:**
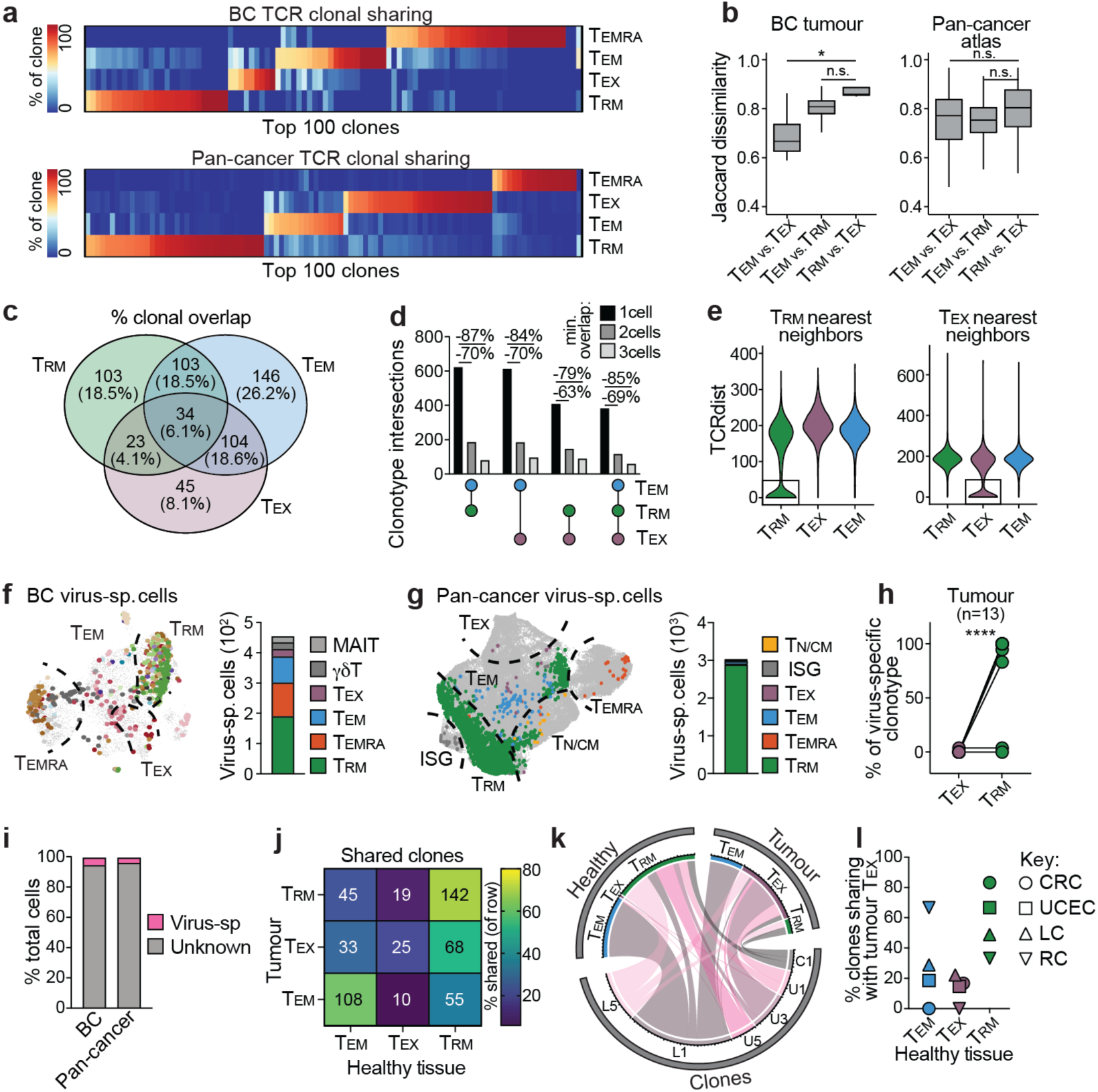
Tumour-associated T_RM_ and T_EX_ cells are clonally distinct with discrete specificities. **a,** Relative frequency of the top 100 expanded TCR clones within each metacluster from the BC dataset (top) or pan-cancer atlas (bottom)^28^. **b,** Jaccard dissimilarity index scores (1 – Jaccard index) for expanded (minimum 2 cells) tumour-derived T cell clones showing dissimilarity between respective populations from BC dataset or pan-cancer atlas, analysed by t-test. **c,** Venn diagram indicating clonal overlap between expanded T_RM_, T_EM_, or T_EX_ clonotypes from BC dataset. **d,** UpSet plot showing the overlap among T_EM_, T_RM_, and T_EX_ cell types from pooled BC, liver, and pan-cancer datasets, with minimum overlap cutoffs of 1, 2, or 3 cells. **e**, Violin plots representing TCRdist analysis of pooled BC, liver, and pan-cancer datasets, showing the distance between T_RM_ or T_EX_ with respective subsets, where lower values indicate greater similarity. **f-g,** Distribution, and enumeration of cells expressing virus-specific TCRs as determined by VDJdb^35–37^ within the BC dataset (**f**) and pan-cancer dataset (**g**). Individual dots indicate virus-specific cells, uniquely coloured by clones (**f**) or phenotype (**g**). **h,** Frequency of virus-specific cells within each clonotype that adopt T_RM_ or T_EX_ cell phenotypes, analysed using a chi-squared test to determine the relative frequency of a virus-specific T cell adopting each phenotype. **i,** frequency of virus-specific T cells of total CD8^+^ T cells in respective datasets. **j-l,** Clonal sharing between tumour and healthy tissue-derived CD8^+^ T cells^40^. **j,** Clonal sharing between subsets detected across tumour vs healthy tissue-derived CD8^+^ T cells, filtered on clones identified in both tissues. Heatmap scaled by % sharing in each row, numbers indicate shared clones pooled from CRC (colorectal), UCEC (uterine corpus endometrial), LC (lung) and RC (renal) cancers^40^. **k,** Circos plot indicating selected clones from distinct cancers, and number of cells from each clonotype occupying each tissue and phenotype. **l**, summary of clonal sharing between tumour T_EX_ and respective phenotype in healthy tissue, split by cancer. n.s. p>0.05, *p<0.05, ****p<0.0001.

Structurally similar TCRs are predicted to recognise similar epitopes. Using TCRdist^33,34^, we identified clear structural segregation between T_RM_ and T_EX_ cell TCRs, with significant structural similarity predicted only among cells of the same phenotype. This finding suggests that T_RM_ and T_EX_ populations possess distinct epitope specificities (**Fig. 4e**). To investigate further, we integrated TCR sequences and HLA allele expression with VDJdb^35–37^ to predict viral reactivity of CD8^+^ T cells in both our BC dataset and the pan-cancer atlas. Predicted virus-specific clonotypes were predominantly associated with a T_RM_ phenotype, while virus-specific T_EX_ cells were exceedingly rare (**Fig. 4f-h**). Interestingly, the propensity to adopt a T_RM_ cell phenotype varied depending on viral specificity. Predicted influenza A-specific cells most frequently exhibited a T_RM_ phenotype, while EBV-specific T_RM_-like cells were rarely identified (Extended Data Fig. 6b-c).

Critically, only a very small frequency of the total CD8^+^ T cells in both datasets was predicted to be virus-specific **(Fig. 4i)**, yet these data indicate that tumour-antigen-independent cells are enriched for the T_RM_ gene signature. In contrast, previous studies have shown that tumour-antigen-specific cells express markers such as CD39^38,39^, which are characteristic of T_EX_ cells in our classification. Based on this, we hypothesised that if tumour-derived clones expressing the T_EX_ gene signature are also found in healthy tissue, where tumour antigen is presumably absent, they may instead show enrichment of the T_RM_ gene signature.

Although clonal sharing between tumour and healthy-tissue-derived cells in our BC and pan-cancer datasets was limited, likely due to inadequate sampling, we leveraged a dataset in which T cell clones were identified in both tumour and adjacent healthy tissue^40^. We extracted CD8^+^ T cell clones shared across sites and annotated them based on our T_RM_ and T_EX_ gene signatures. Consistent with our hypothesis, there was significant clonal sharing between tumour-derived T_EX_ and healthy tissue-derived T_RM_ cells **(Fig. 4j-k, Extended Data Fig. 6d-e)**. Indeed, when clonal overlap was observed, tumour-derived T_EX_ clonotypes were more frequently shared with healthy T_RM_ than with other T cell types **(Fig. 4l)**. Together, these data suggest that the T_RM_ gene signature is enriched in predicted tumour-antigen-independent memory T cells, and tumour-specific cells in healthy non-cancerous tissues.

### Antigen drives the distinction between T_RM_ and T_EX_ cells in tumours

Given that putative tumour-antigen-specific T cells preferentially adopt a T_RM_ phenotype in surrounding tissues where tumour antigen may be absent, we hypothesised that tumour-antigen reactivity drives the divergence between tumour-associated T_RM_ and T_EX_ cells. To test whether tumour-specific T_RM_ can form in tumours, we utilised an orthotopic murine BC (AT3) model engineered to express the model antigen ovalbumin (OVA). High-dimensional flow cytometry of tumour-infiltrating T cells (TILs) revealed 4 clusters enriched for tumour-specific OT-I T cells expressing CD69 and PD-1, and differing in markers such as CD39, CD103, Tcf1, Tox and Tim3 **(Fig. 5a-f, Extended Data Fig. 7a)**. We annotated these subsets based on phenotypic and functional characteristics. First, we defined resident cells as non-migratory. In lieu of definitive migration assays (e.g., parabiosis), we used indirect indicators of tissue residency, including intravenous (IV) labelling and FTY720 treatment. Over 90% of TILs were IV-negative, and CD69^+^ TILs showed reduced IV-staining relative to CD69^-^ TILs **(Extended Data Fig. 7b)**. TIL frequencies were also unaffected by FTY720 treatment **(Extended Data Fig. 7c)**, supporting the notion that the majority of TIL are resident.

**Figure 5:**
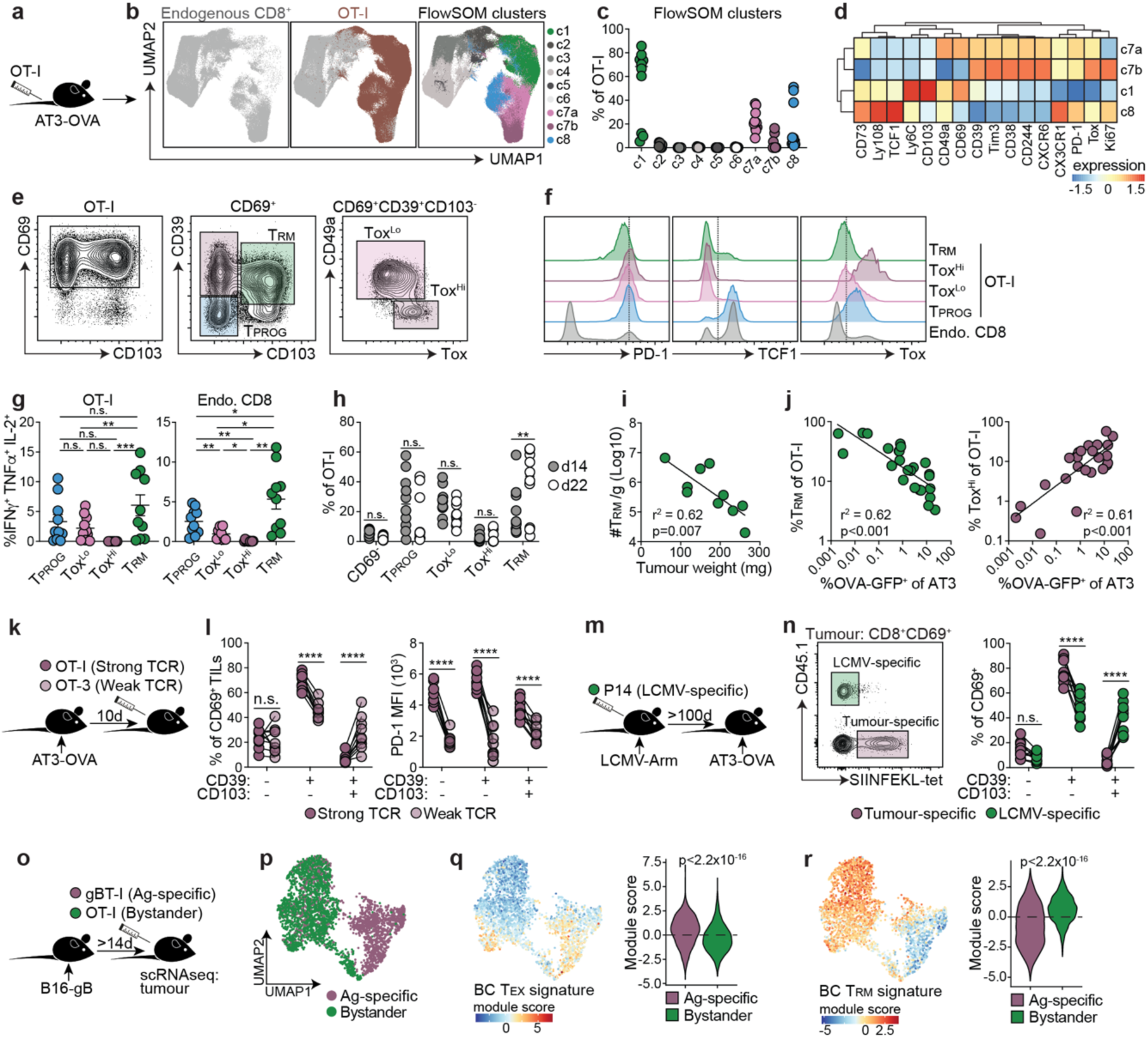
Low-avidity and bystander CD8^+^ T cells preferentially adopt a T_RM_ phenotype in tumours. **a-f,** Mice received 1×10⁴ naïve OT-I T cells and were challenged with AT3-OVA; tumour-infiltrating T cells were analysed on day 22. **a,** Experimental schematic. **b,** UMAP of CD8⁺ TILs (defined by markers in Extended Data Fig. 7a) showing OT-I cells and FlowSOM clusters; Cluster 7 was stratified by Tox expression (into c7a (Tox^Lo^) and c7b (Tox^Hi^)). **c,** Distribution of OT-I cells across FlowSOM clusters. **d,** Relative marker expression by cluster. **e-f,** Representative gating **(e)** and marker expression **(f)** from tumour-derived OT-I cells, Endo. CD8: endogenous CD8⁺ T cells. **g,** Cytokine expression post-PMA/ionomycin restimulation. **h,** OT-I frequencies at d 14 and 22. **i,** T_RM_ cell number per gram tumour vs. tumour mass at day 14. **j,** Proportion of T_RM_ and Tox^Hi^ OT-I vs. %GFP-OVA^+^ AT3 at d 22. **k-l,** Effector OT-I (strong TCR signaling) and OT-3 (weak TCR signaling) T cells were co-transferred into tumour-bearing mice (day 10); intratumoural T cells were analysed on day 28. **k,** Schematic. **l,** Frequency of CD69⁺ cells and PD-1 expression. **m-n,** LCMV-immune mice (receiving 5×10⁴ P14 prior to infection) were challenged with AT3-OVA >100d post-LCMV infection; intratumoural T cells analysed on day 23 post-tumour inoculation. **m,** Schematic. **n,** Frequencies of CD45.1⁺ LCMV-specific (P14) vs. SIINFEKL-tetramer⁺ tumour-specific cells among CD8⁺CD69⁺ TILs. **o-r,** scRNA-seq of bystander (OT-I) and Ag-specific (gBT-I) CD8⁺ T cells co-transferred into mice with B16-gB melanoma tumours. **o,** Schematic. **p,** UMAP of sorted cells coloured by transgenic T cell. **q-r,** Overlay of BC T_EX_ (q) and T_RM_ (r) gene signatures. p-values from Wilcoxon signed-rank test. Statistics: **b-j,l,** pooled from 2 independent experiments; **n,** representative of 2 experiments; **b-f,** representative of >8 total experiments, minimum N=5 mice/group/experiment. **g:** one-way repeated-measures ANOVA with Tukey post-test; **h:** two-way ANOVA with Sidak test; **i,j:** r^2^ indicates fit of linear regression line and p-value indicates slope departs from zero. **l,n:** two-way repeated-measures ANOVA with Sidak test. **l,n:** connected points from individual mice. n.s., p > 0.05; *p<0.05, **p < 0.01; ***p < 0.001; ****p < 0.0001.

Second, we considered cells exhausted if they exhibited functional impairment, and memory cells as those capable of persisting without ongoing antigen stimulation. Accordingly, CD39^+^ CD103^+^ cells were annotated as T_RM_ cells based on superior functionality, while CD39^+^ CD103^-^ Tox^Hi^ cells represented the most dysfunctional, bona fide exhausted population **(Fig. 5g, Extended Data Fig. 7d-e)**. A separate CD69^+^ CD39^-^ CD103^-^ population expressed Tcf1, PD-1, and Tox, consistent with progenitor exhausted T cells (T_PROG_), which were selectively depleted when Tcf7 expression was ablated **(Fig. 5e-f, Extended Data Fig. 8a-d)**. Strikingly, we found that the number of T_RM_ phenotype cells inversely correlated with tumour size at early timepoints (d14) **(Fig. 5i)**, and acquisition of the T_RM_ phenotype correlated with the loss of OVA-GFP expression from tumour cells over time (d23) **(Fig. 5j)**. In contrast, Tox^Hi^ exhausted cells were enriched in tumours that maintained high tumour-antigen load, consistent with the role of antigen in driving T cell exhaustion **(Fig. 5j)**. Inducible TCR depletion in tumour-specific T cells increased both the frequency and number of CD103^+^ T cells (**Extended Data Fig. 8e-h)**, further supporting that CD103 expression marks memory T cells in this model, and that tumour-specific T_RM_ cells preferentially form or persist when cognate antigen is absent, consistent with classical T cell memory paradigms.

Given that the presence or absence of intratumoural TCR signalling appeared to influence T_EX_ versus T_RM_ cell fate, we next assessed whether reducing TCR signal strength would also influence this distinction. Indeed, OT-3 T cells, which have reduced TCR signalling towards OVA-peptide compared to OT-I T cells, displayed increased CD103 and decreased PD-1 expression (**Fig. 5k-l)**, indicating that low-avidity TCR signalling promotes T_RM_ cell differentiation within tumours. Consistent with the T_RM_ phenotype of virus-specific T cells in human tumours, virus-specific memory T cells generated >100 days after LCMV infection infiltrated AT3-OVA tumours and adopted a T_RM_ phenotype (**Fig. 5m, n**). Furthermore, transfer of tumour-specific (OT-I) and non-specific bystander (gBT-I) effector T cells into tumour-bearing mice showed that bystander T cells more readily acquired a T_RM_ phenotype (**Extended Data Fig. 8i-j**). These bystander T cells expressed significantly lower levels of T_EX_-related proteins such as PD-1, underscoring phenotypic differences between tumour-specific and bystander-derived T_RM_ populations (**Extended Data Fig. 8j**). The development of intratumoural bystander T_RM_ cells required intrinsic TGFβ-signaling **(Extended Data Fig. 8k-l),** and their efficient tumour entry was dependent on CXCR3 and CXCR6 expression **(Extended Data Fig. 8m-n)**.

Given the apparent correlation between antigen presence and T_RM_ cell biasing, we speculated that the distinct transcriptional T_EX_ and T_RM_ cell signatures derived from patient samples above may similarly correlate with tumour-antigen reactivity. To assess this, we performed scRNAseq on tumour-specific (gBT-I) and bystander (OT-I) T cells derived from B16-gB tumours and observed discrete clustering of these populations (**Fig. 5o,p**). Notably, tumour-specific T cells displayed enrichment of the human BC T_EX_ gene signature (**Fig. 5q**), whereas bystander T cells exhibited high expression of the BC T_RM_ gene signature (**Fig. 5r**). Consistently, bystander T cells from E0771 BC tumours^16^ expressed the T_RM_ gene signature and tumour-specific T cells expressed the T_EX_ gene signature (**Extended Data Fig. 8o-q**). Thus, the relative presence or absence of intratumoural TCR signalling drives the distinct transcriptional profiles of tumour-associated T_RM_ and T_EX_ cells. Altogether, these data indicate that strong and persistent TCR signalling is antithetical to T_RM_ cell development, suggesting that bona fide T_RM_ cells within human tumours are likely tumour-agnostic bystanders, low-affinity tumour-specific T cells, or T cells not in contact with their cognate antigen.

### Tumour antigen drives T_RM_ cells towards a T_EX_ cell fate

The observation that T_RM_ and T_EX_ cells are clonally distinct in human tumours likely reflects intrinsic differences in antigen reactivity, rather than indicating that these populations derive from separate precursor cells. Notably, this clonal distinction does not preclude the presence of tumour-specific T_RM_ cells, nor the possibility of clonal overlap with T_EX_ cells. Indeed, our analyses suggested that such clonal overlap could occur when tumour-specific T_RM_ cells are present in tumours or in tissues where cognate antigen is presumably absent. As such, these results did not resolve whether tumour-specific T_RM_ and T_EX_ cells arise from a shared progenitor or from distinct developmental lineages. To address this, we employed the SPLINTR barcoding system^41^ to introduce unique barcodes into mono-specific naïve or effector OT-I T cells before adoptive transfer into AT3-OVA tumour-bearing mice (**Fig. 6a**). To generate naïve barcoded T cells, we transduced hematopoietic stem cells from OT-I donor mice with SPLINTR-encoding lentivirus and performed intra-thymic injections into sub-lethally irradiated recipients. 8 weeks later, we pooled naïve T cells from 20 chimeric donor mice to increase barcode diversity and transferred either 2×10^3^ or 1×10^4^ naïve OT-I T cells into recipient mice subsequently inoculated with AT3-OVA. 24 days after tumour inoculation, we sorted SPLINTR-barcoded OT-I T cells from spleens and tumour populations, including CD69^-^, T_PROG_, T_RM_, and the heterogeneous CD39^+^CD103^-^ population we refer to as “T_EX_-like”. Barcode distribution was assessed by DNA sequencing.

**Figure 6:**
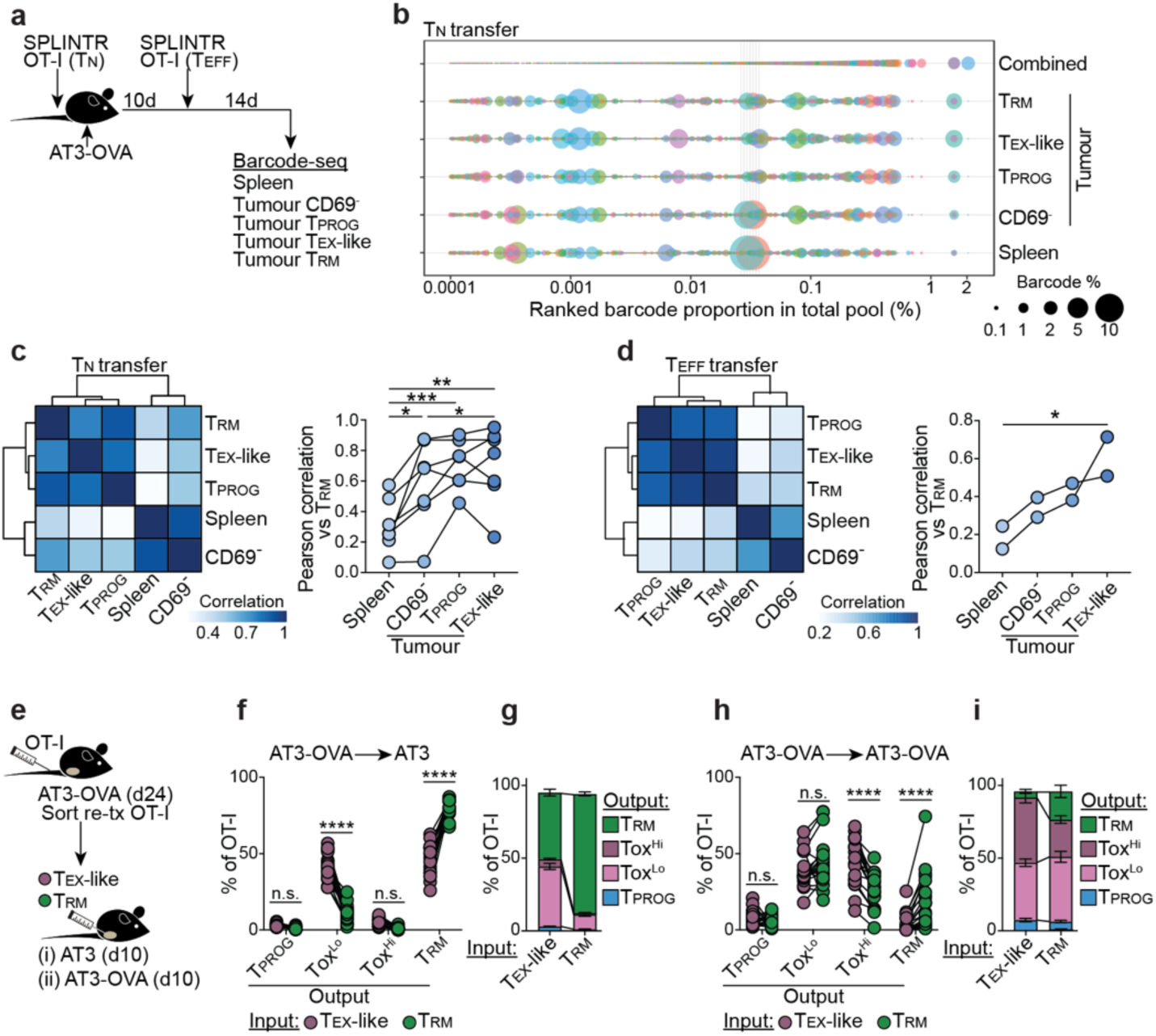
Tumour-associated T_RM_ cells can be driven towards exhaustion. **a-d,** SPLINTR barcode-seq of naïve (T_N_) or effector (T_EFF_) OT-I T cells transferred into mice AT3-OVA tumour-bearing mice. **a,** Schematic. TIL populations as defined in Fig. 5e. T_EX_-like population includes Tox^Lo^ and Tox^Hi^ populations. **b,** Barcode representation in a T_N_ transferred mouse sorted by total pooled barcode where bubble size reflects clone proportion in sample. **c-d,** Pearson correlation of barcodes identified in respective populations from representative mouse (heatmaps) and all repeats compared to T_RM_ population (line-plot) from T_N_ transfer (c) or T_EFF_ transfer (d). **e-i,** Sort-retransfer of respective OT-I populations from AT3-OVA mice. **e,** Schematic. Sorted cells were co-transferred intratumourally into recipient mice. **f-g**, % of populations isolated from AT3 bearing recipient mice, split by input cell phenotype, and output phenotype in recipient mouse, showing independent mice (f) and summary (g). **h-i**, % of populations isolated from AT3-OVA bearing recipient mice, split by input cell phenotype, and output phenotype in recipient mouse, showing independent mice (h) and summary (i). Statistics: **b-c,** representative of 8-independent biological replicates from 3 experiments; 1-way repeated measures ANOVA with Tukey post-test. **d,** representative of 2-independent biological repeats each with technical replicates (not shown); 1-way repeated-measures ANOVA with test for linear trend. **f-i,** pooled from 2-independent experiments, N=19 total mice; 2-way repeated measures ANOVA with Sidak post-test n.s. p>0.05, *p<0.05, **p<0.01, ***p<0.001, ****p<0.0001.

Minimal barcode sharing was observed between mice receiving 2×10^3^ OT-I T cells, and sharing was only detectable in mice that received 1×10^4^ cells **(Extended Data Fig. 9a)**, indicating that most barcodes were unique and minimising the possibility of PCR artifacts. To avoid the use of highly duplicated barcodes in the original naïve pool, we excluded any barcodes identified across multiple mice before assessing barcode sharing across splenic and tumour populations within individual mice. These analyses showed that antigen-specific, tumour-derived T_PROG_, T_EX_-like, and T_RM_ cells were more similar to each other, and distinct from CD69^-^ and spleen-derived populations **(Fig. 6b-c)**. We conducted a parallel experiment using effector OT-I T cells transduced with SPLINTR-encoding retrovirus, transferred into AT3-OVA-bearing mice. Barcode diversity was again determined across spleen and tumour populations, confirming that barcoded OT-I library pools were unique (**Extended Data Fig. 9b**). As per the naïve T cell experiment, barcode distribution reinforced that tumour-derived T_PROG_, T_EX_-like, and T_RM_ cells displayed a high degree of barcode overlap and were distinct from CD69^-^ and spleen-derived populations (**Fig. 6d, Extended Data Fig. 9c**). Altogether, these data indicate that tumour-specific T_RM_ cells do not arise from a distinct T cell lineage but instead share common progenitors with other intratumoural populations such as T_EX_-like and T_PROG_ cells.

The developmental association between each cell type left unresolved whether tumour-antigen-specific T_RM_ cells that develop after antigen loss or following spatial segregation from cognate antigen can transition into T_EX_ cells upon antigen re-encounter. To investigate this, we isolated intratumoural OT-I T cells exhibiting T_PROG_, T_EX_-like, or T_RM_ phenotypes from AT3-OVA tumours and re-transferred them intravenously into secondary recipient mice bearing AT3-OVA tumours (**Extended Data Fig. 9d**). Among these populations, T_EX_-like OT-I T cells were significantly less efficient at repopulating tumours compared to T_PROG_ or T_RM_ cells (**Extended Data Fig. 9e-f**). Due to the relatively low recovery of transferred cells, we performed composite protein expression profiling from each transferred group by experiment, and compared these profiles to T_PROG_, T_EX_-like, and T_RM_ OT-I T cells isolated from concurrently analysed primary tumours (**Extended Data Fig. 9g-h**). This analysis revealed that all re-transferred populations in secondary tumours most closely resembled the T_EX_-like phenotype observed in primary tumours.

To determine whether differences in trafficking influenced the ability of cells to repopulate tumours and adopt distinct phenotypes, we sorted congenically distinct T_EX_-like and T_RM_ OT-I T cells from primary AT3-OVA tumours and co-transferred them directly into secondary tumours that either expressed or lacked OVA **(Fig. 6e)**. Notably, direct intratumoural transfer eliminated the repopulation advantage previously observed for T_RM_ cells, potentially reflecting their enhanced ability to repopulate distal sites following intravenous transfer **(Extended Data Fig. 9i)**. In the absence of cognate antigen, transferred T_RM_ cells largely retained their phenotype, whereas approximately 50% of the heterogenous T_EX_-like population (input, comprising Tox^Lo^ and Tox^Hi^) could adopt the T_RM_ phenotype **(Fig. 6f-g)**. Strikingly, the dysfunctional Tox^Hi^ population was absent within AT3 tumours lacking antigen, consistent with earlier data showing these cells are sustained in the presence of antigen **(Fig. 6f-g)**. Conversely, when the same populations were transferred into AT3-OVA tumours, a fraction of T_RM_ cells maintained their phenotype; however, the majority transitioned to a T_EX_-like state, including a significant proportion acquiring the Tox^Hi^ phenotype **(Fig. 6h-i)**. Thus, T_RM_ cells can be driven towards terminal exhaustion within the antigen-rich environment of secondary tumours. Overall, these data indicate that, unlike non-tumour reactive bystander T_RM_ cells, tumour-specific T_RM_ and T_EX_ populations can arise from a common origin, with tumour-specific T_RM_ cells having the capacity to be driven towards exhaustion upon chronic antigen re-encounter.

## Discussion

Our study for the first time reconciles two critical T cell subsets associated with tumour control, namely CD8^+^ T_RM_ and T_EX_ cells. We show that these subsets are distinct T cell populations that have been conflated in the literature. This conflation has arisen due to T_EX_ cells engaging a residency program that relies on the same transcriptional machinery used by T_RM_ cells to inhibit tissue egress. Thus, T_RM_ gene signatures developed without T_EX_ cell consideration cannot disentangle these two populations. This study resolves this issue by establishing broad gene signatures that reliably distinguish these subsets across various human tumours.

The deconvolution of tumour-associated T_RM_ and T_EX_ cells highlights their potential to play distinct roles in anti-tumour immunity and their differential impact on ICB responses. High T_EX_ gene signature expression in BC correlates with positive ICB outcomes reflecting the therapy’s aim to reinvigorate T_EX_ cells through blockade of inhibitory receptors like PD-1 and CTLA4 which are expressed at much higher levels in T_EX_ than T_RM_ cells. The association of high TMB with increased T_EX_ cell frequency suggests that increased novel tumour epitopes enable T_EX_ cell development. T_EX_ cells, as defined in this study, consistently co-express CD39 and CD103, markers known to enrich tumour-specific CD8^+^ T cells^38,39,42^. High TMB is generally associated with better ICB responses, including BC subtypes such as triple-negative breast cancer (TNBC), where TMB predicts favourable outcomes to therapies like pembrolizumab^43–46^. The observed link between high T_EX_ gene signature scores, TMB, and ICB efficacy underscores the critical role of T_EX_ cells in tumour control.

We found that most tumour-associated T_RM_ cells identified in humans are clonally and developmentally distinct from T_EX_ cells. Tumour-antigen agnostic T cells, including T cells specific for previous infections, can adopt a residency phenotype within tumours. Consistently, we show that TCR signalling is antithetical to bona fide T_RM_ cell formation in tumours, aligning with the conceptual notion that immunological memory can only develop following clearance of cognate antigen. However, we also reveal that tumour-antigen-specific T_RM_ cells can develop in settings of reduced TCR signaling and diminished antigen sensing. Tumour-specific T_RM_ cells likely exist in lower numbers compared to tumour-specific T_EX_ cells since it is expected that they have encountered less antigen, and therefore had significantly reduced proliferative bursts, accounting for the limited clonal sharing that we observe between T_RM_ and T_EX_ cells in human tumours.

While T_RM_ gene signatures are associated with overall survival in BC patients, the mechanisms underlying T_RM_-mediated tumour control remain unclear. It is possible that the accumulation of these T_RM_ cells may coincide with other features of tumour control, such as tumour antigen clearance. However, our analyses suggest that tumour-specific T_RM_ cells exist in healthy tissues surrounding tumours, raising the possibility that they contribute to long-term immune surveillance and protection against tumour recurrence. Thus, the clinical benefit of tumour-specific T_RM_ cells may lie in their ability to maintain equilibrium with residual cancerous cells, thereby preventing tumour recurrence, or in their potential prophylactic use following vaccination^47^. Given that these cells develop after clearance of their cognate antigen and have an increased ability to traffic and repopulate distal sites, the presence of T_RM_ cells in human tumours may reflect an effective immune response against tumour antigen that can lead to lasting protection both at the primary tumour location, and potential sites of metastasis.

Finally, significant populations of bona fide, non-exhausted, tumour-agnostic T_RM_ cells can be identified within diverse tumours. These cells maintain superior functionality in humans and murine models but are not targeted by current immunotherapies. Thus, approaches to activate bystander T_RM_ cells via TCR-independent pathways or administration of viral peptides^39,48,49^, or to redistribute functional tumour-specific T_RM_ cells, combined with current T_EX_ cell-targeted ICB therapies, could raise the ceiling of effective anti-tumour responses.

## Extended Data

**Extended Data Figure 1:**
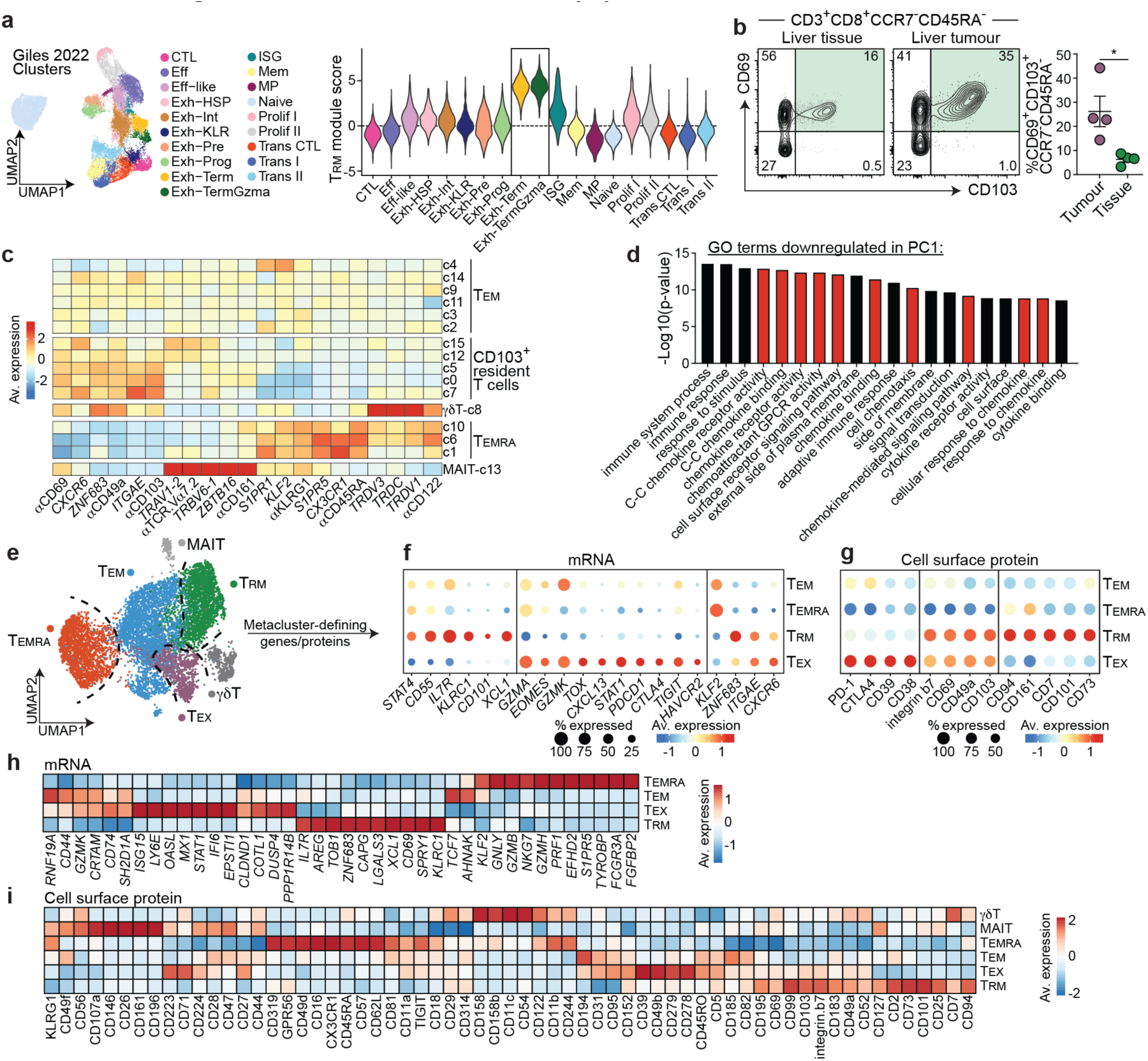
Identification of CD8^+^ T cell populations in tumours. **a,** T_RM_ gene signature expression on LCMV-specific CD8^+^ T cell clusters from scRNA-seq of the respective dataset^27^ (Related to Fig. 1b-d). **b**, Flow cytometry of CD8^+^ T cells isolated from liver tumours and liver tissue from N=4 colorectal cancer patients with liver metastases. Representative plots and summary data for CD69^+^CD103^+^CCR7^-^CD45RA^-^ CD8^+^ T cells, analysed by Mann-Whitney test, *p<0.05. **c,** Expression of respective lineage-defining genes and cell surface proteins from BC-CITEseq data (Fig. 1i-j) displayed on heatmap by cluster. **d,** pathway enrichment of downregulated loading genes associated with PC1 from PCA plot in Fig. 1n. Red bars indicate pathways related to chemokine receptor signaling and trafficking **e,** Schematic showing metacluster annotations. **f-g**, Expression of selected shared and distinct T_RM_ and T_EX_ genes (**f**) or cell surface proteins (**g**) across metaclusters from BC dataset. **h-i**, heatmaps showing top DE genes (**h**) and cell surface proteins (**i**) from BC dataset.

**Extended Data Figure 2:**
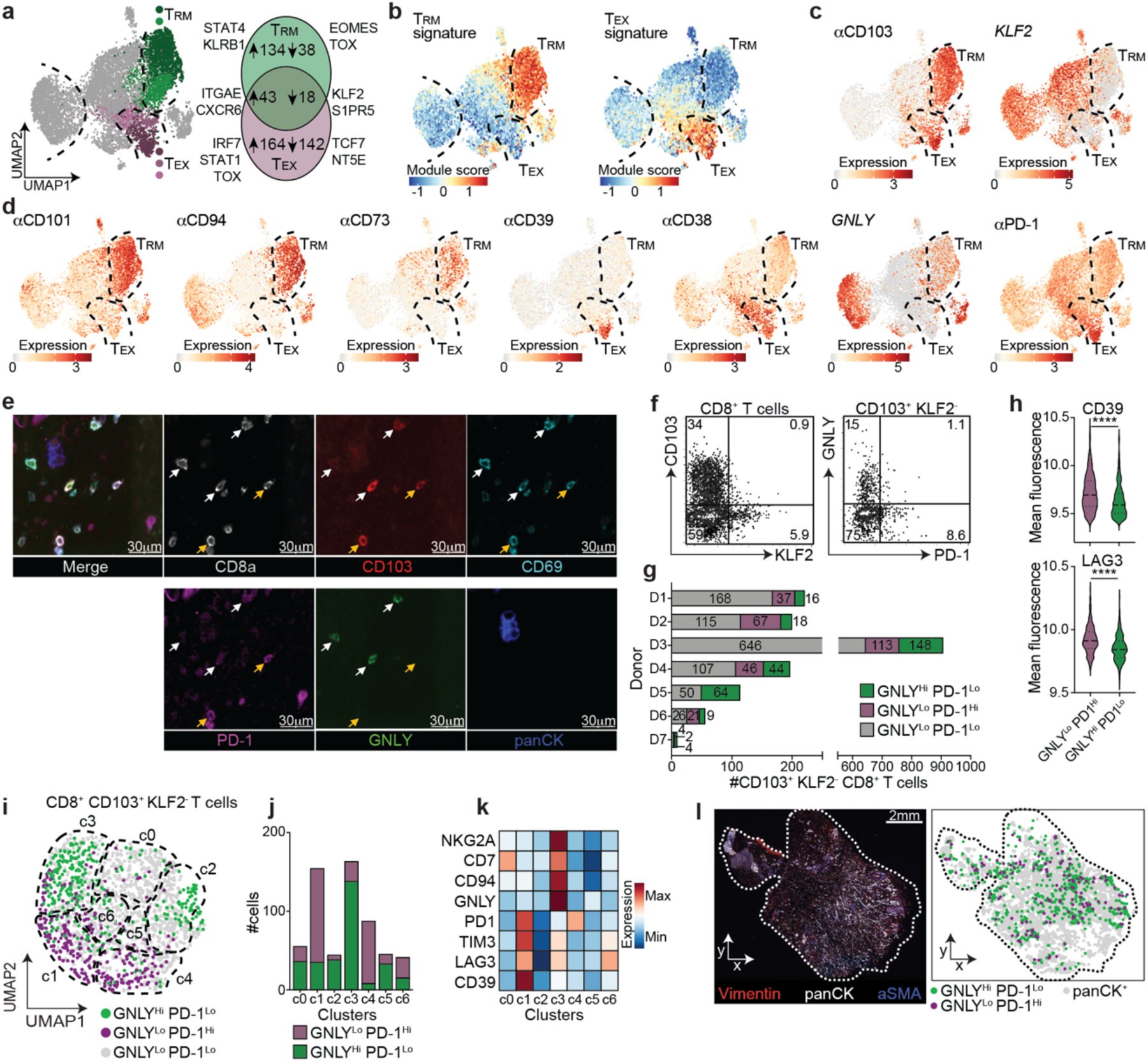
Lineage-defining genes and proteins enable *in situ* localisation of T_RM_ and T_EX_ cells. **a,** T_RM_ and T_EX_ signature genes. Shared genes indicate genes are up (or down)-regulated in the T_RM_ vs all *and* T_EX_ vs. all comparisons (Fig. 2a). T_RM_ (**b**) and T_EX_ (**c**) gene signature module scores displayed on BC UMAPs for summary module score data (Fig. 2b-c). **c-d**, Expression of respective genes and cell surface proteins displayed on BC UMAPs. **e-l,** CycIF imaging of BC tumours related to Fig. 2f-h. **e**, representative image of BC tumour using CycIF approach showing respective stains. **f**, Histocytometric gating strategy showing CD8^+^ T cells (CD3^+^ CD8^+^ CD4^-^) concatenated from CycIF images from 7 donors showing CD103, KLF2, GNLY, and PD-1 expression on respective populations. **g,** number of CD8^+^ CD103^+^ KLF2^-^ T cells identified in each slide from respective donors, broken down by GNLY and PD-1 expression based on gates in (f). **h**, Expression of CD39 and LAG3 proteins on respective populations, pooled from 7 donors. Mann-Whitney test ****p<0.001. **i**, UMAP projection and clustering of CD8^+^ CD103^+^ KLF2^-^ T cells, coloured by gates defined in (f). **j**, number of cells from respective gates identified within each cluster. **k**, relative expression of respective proteins in each cluster. **l,** representative image of tumour section showing panCK and aSMA expression, and relative location of respective annotated cell types.

**Extended Data Figure 3:**
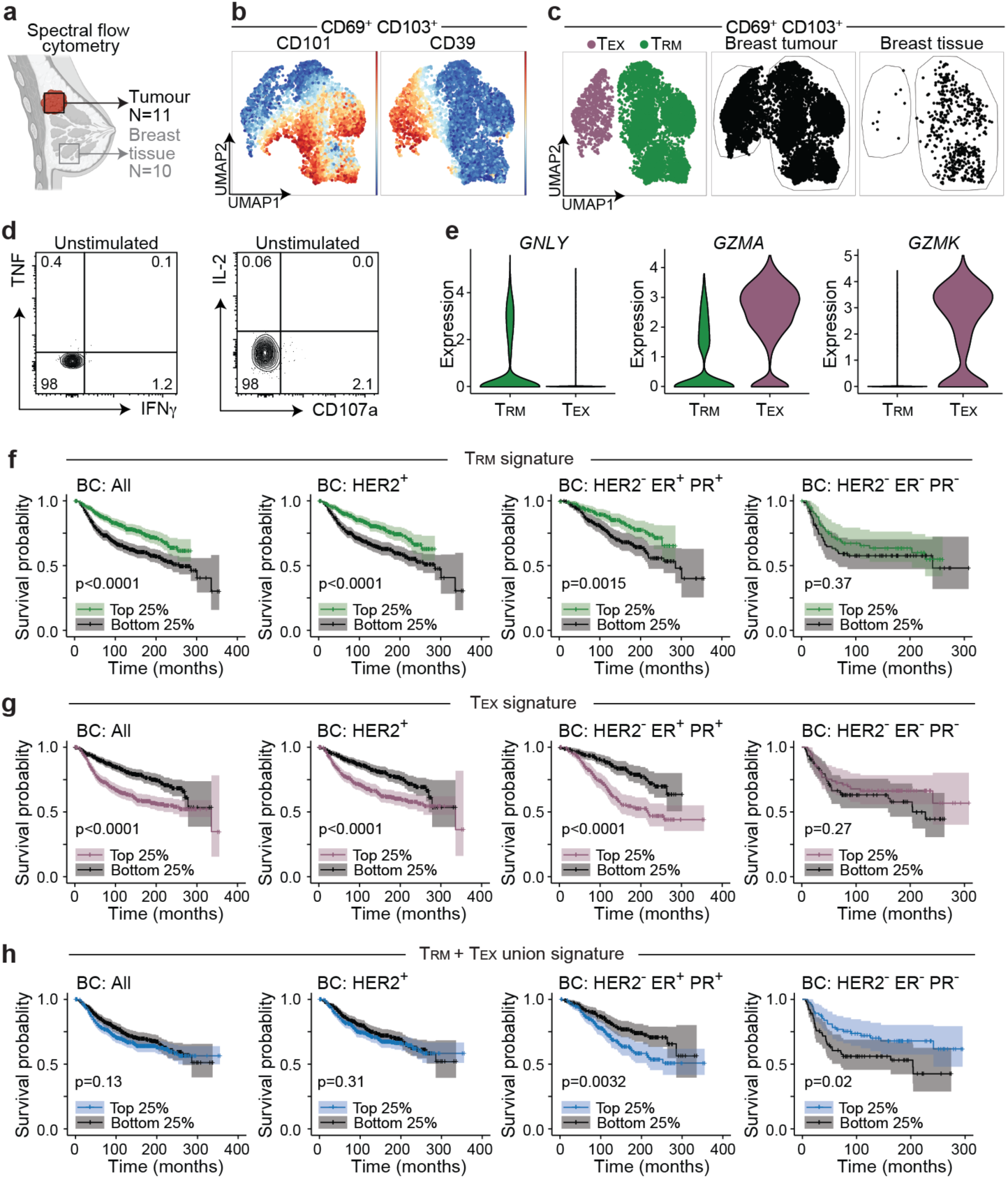
Functional assessment and survival associations of BC T_RM_ and T_EX_ cells. **a-e,** Flow cytometry of PMA-ionomycin restimulated BC and breast tissue-derived CD103^+^ resident-phenotype T cells related to Fig. 2d-k. **a**, Schematic. **b-c,** CD69^+^CD103^+^CD45RA^-^CCR7^-^ CD8^+^ T cells represented in UMAP space based on the expression of CD69, CD101, CD39, CD94, CD73, CD103, CD45RA, CCR7, CD38 showing CD101 and CD39 expression (**b**) and annotated T_RM_ and T_EX_ populations and relative contribution by cells derived from BC tumours or tissue (**c**). **d,** Gating of TNF, IFNγ, IL-2, and CD107a for Fig. 2i-k based on unstimulated control. **e,** Expression of granulysin (*GNLY*), granzyme A (*GZMA*) and granzyme K (*GZMK*) on T_RM_ and T_EX_ populations from BC CITEseq dataset. **f-h,** Survival of BC patients from the dataset with the highest (top 25%) T_RM_ (**f**), T_EX_ (**g**), or combined T_RM_ + T_EX_ union (**h**) gene signature score enrichments compared to patients with lowest (bottom 25%) gene enrichment scores and plotted on Kaplan-Meier curves with log-rank test, as in (Fig. 2l), segregated by BC subtypes (as annotated).

**Extended Data Figure 4:**
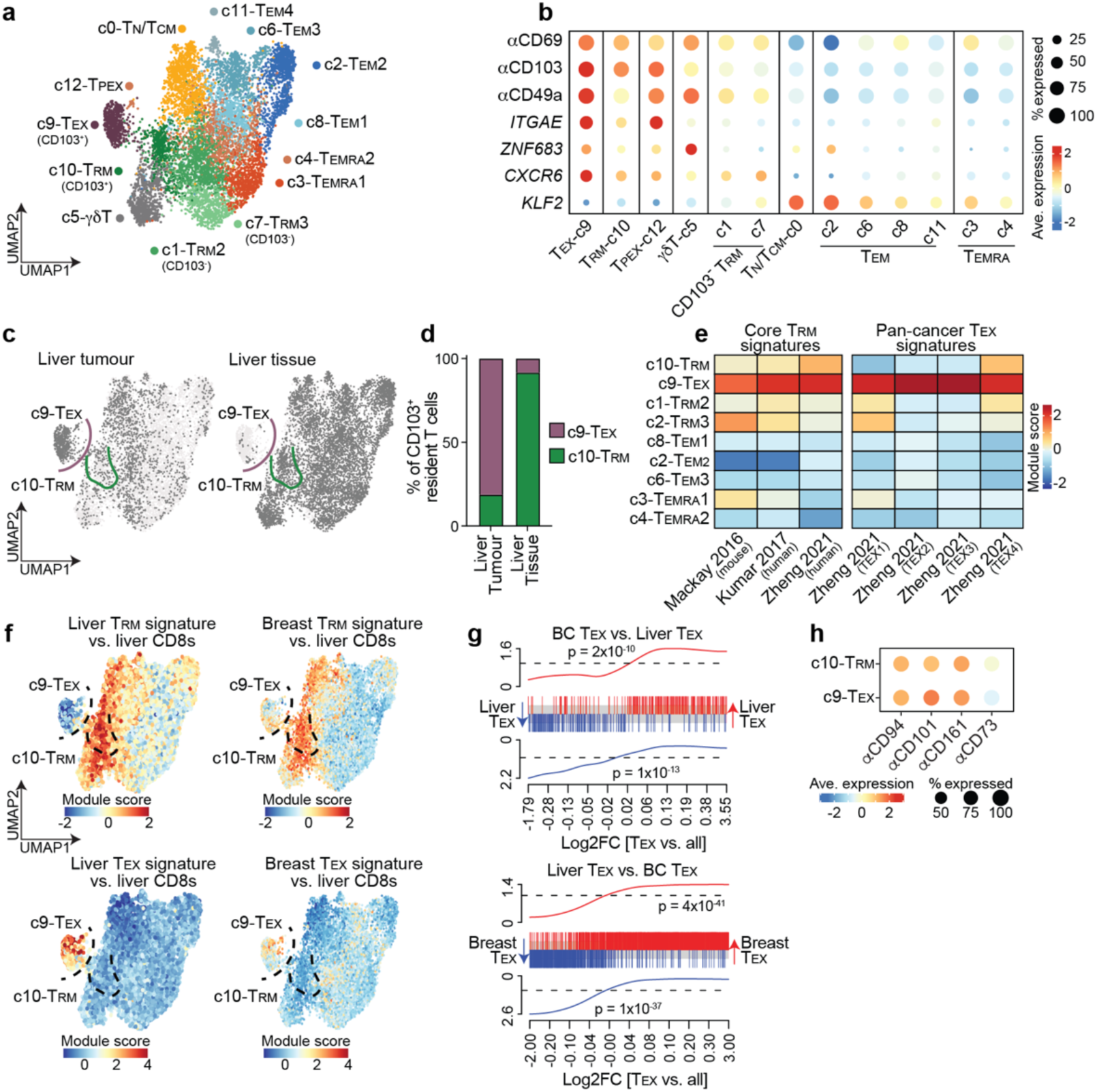
Deconvolution of CD103^+^ T_RM_ and T_EX_ cells in liver metastases. **a-d,** CITEseq of CD3^+^CD8^+^CD4^-^ non-MAIT cells isolated from secondary liver tumours (colorectal cancer patients, N=4) and non-cancerous liver tissue (N=6) as depicted in Fig. 3a. **a,** Data was Harmony integrated, and unified protein and RNA-seq data represented on weighted nearest neighbours UMAP, coloured by clusters that were annotated based on expression of lineage-defining cell surface proteins and genes. **b,** Expression of selected cell surface proteins (αCD69, αCD103,) and genes (*ITGAE, ZNF683, CXCR6, KLF2*) on respective clusters. **c-d,** CD8^+^ T cells segregated by tissue of origin (**c**), and relative cluster composition of CD103^+^ resident T cells (c9+c10) isolated from BC tumours or tissue (**d**). **e,** Average module scores of published T_RM_^8,21,28^ and T_EX_^28^ cell gene signatures by annotated clusters. **f,** Module score overlays of relevant signatures on liver CITEseq dataset. **g,** gene-set enrichment analysis of BC T_EX_ vs liver T_EX_ signatures and vice versa. **h,** Expression of respective cell surface proteins across annotated clusters.

**Extended Data Figure 5:**
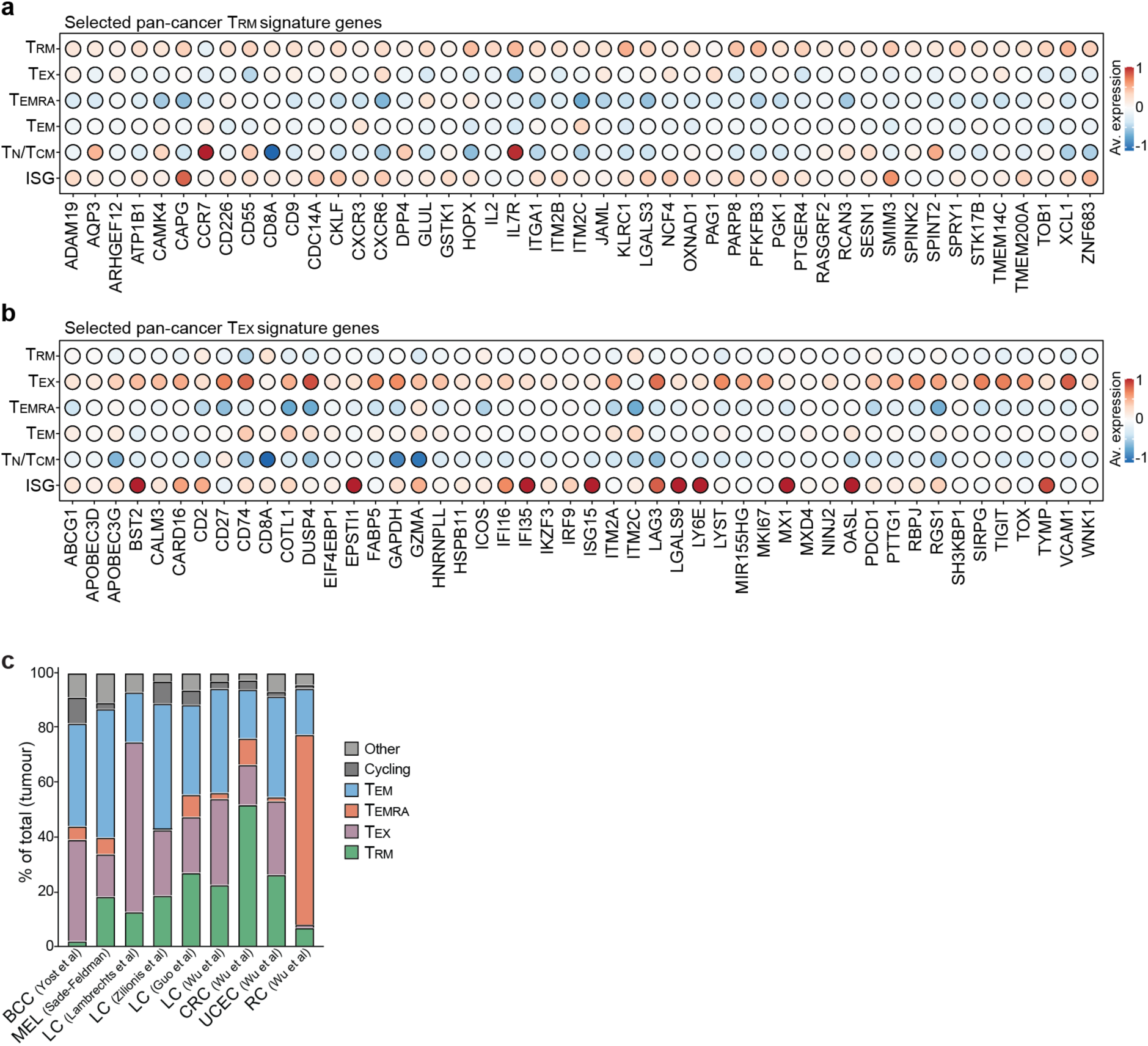
Pan-cancer T_RM_ and T_EX_ signature genes. **a-b**, Relative expression of the top genes from pan-cancer T_RM_ (**a**) and T_EX_ (**b**) gene signatures across respective CD8^+^ T cell populations in the pan-cancer atlas^28^. **c**, Relative frequencies of different CD8^+^ T cell subsets by cancer from respective datasets ^48–53^.

**Extended Data Figure 6:**
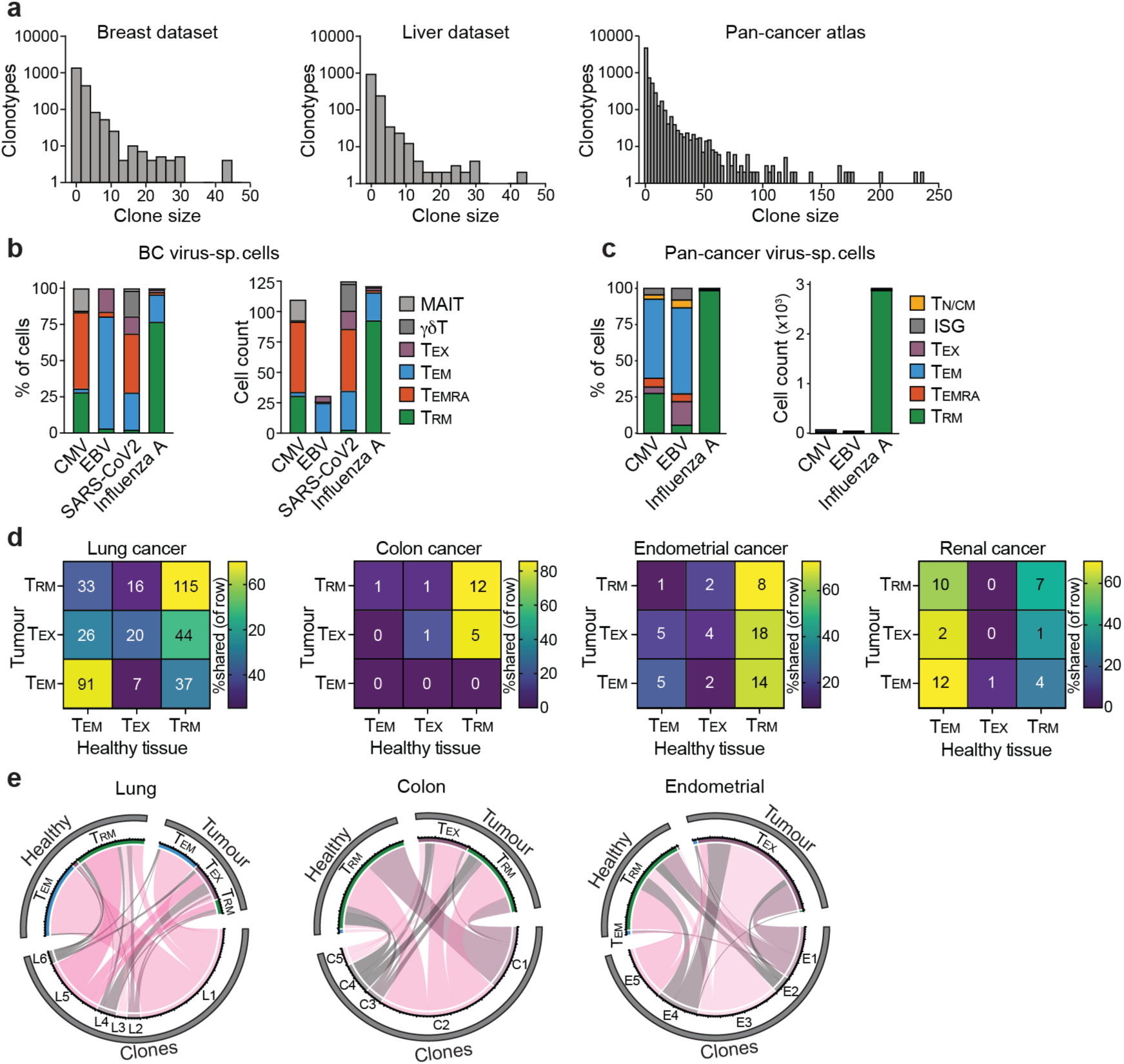
Assessment of virus-specific clones and TCR sharing across tissues. **a,** Clone size distribution across respective datasets. **b-c**, Frequency and count of cells from BC dataset (b) and pan-cancer atlas (c) split by respective virus-specificity. **d-e,** Clonal sharing between tumour and healthy tissue-derived CD8^+^ T cells^40^ related to Fig. 4j-l. Clonal sharing between subsets detected across tumour vs healthy tissue-derived CD8^+^ T cells, filtered on clones identified in both tissues. **d,** Heatmap scaled by % sharing in each row, split by respective cancers^40^. **e,** Circos plots indicating selected clones from respective cancers, and number of cells from each clonotype occupying each tissue and phenotype.

**Extended Data Figure 7:**
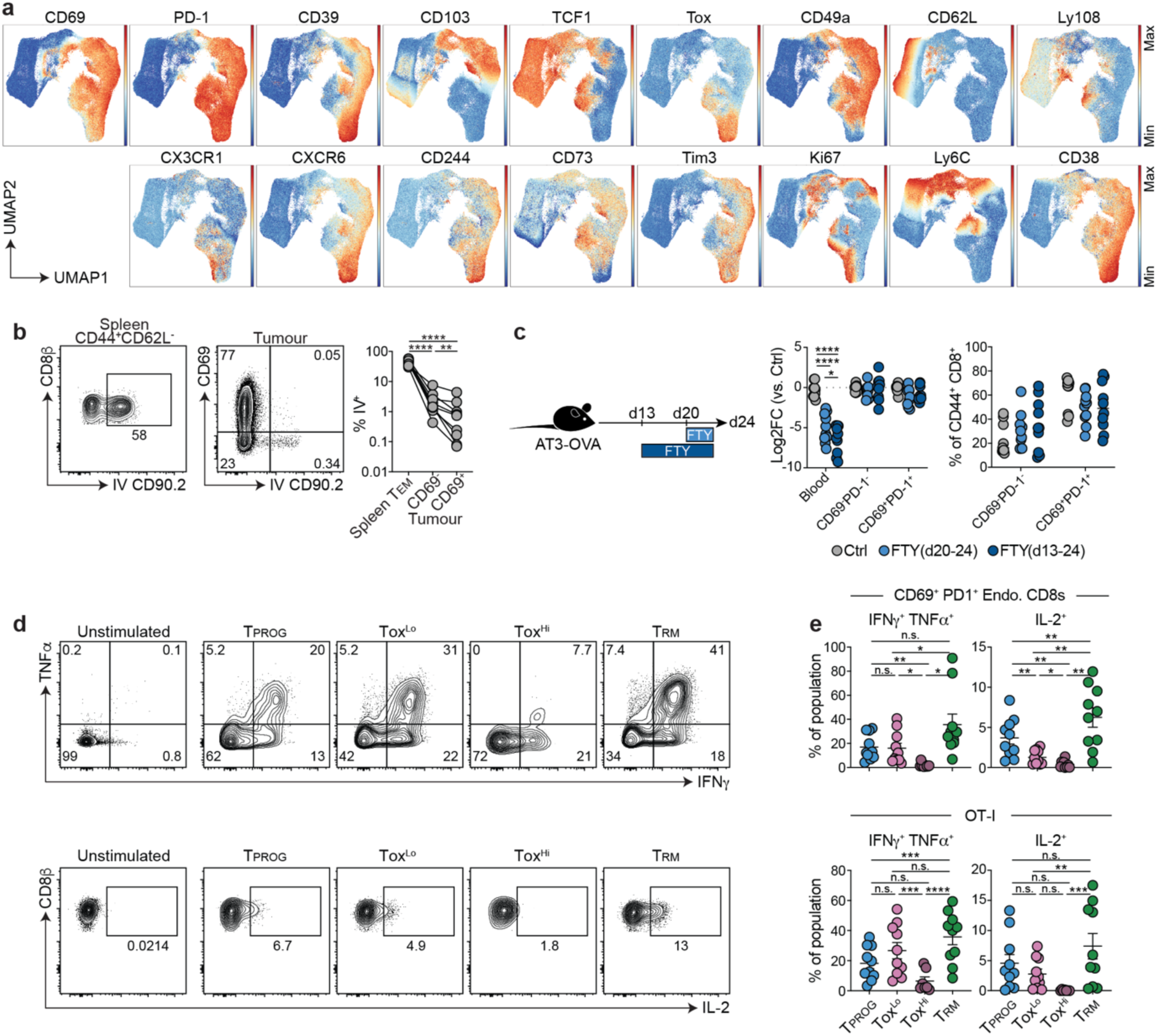
Phenotypic and functional assessment of AT3-OVA CD8^+^ TIL populations. **a**, Expression of respective proteins on CD8^+^ T cells isolated from AT3-OVA tumours projected in UMAP space, related to Fig. 5a-d. **b**, Gating and intravenous (IV) labelling of respective CD8^+^ T cell populations from spleen and tumour at d24 post-inoculation. **c**, FTY720 treatment of AT3-OVA bearing mice showing relative cell number (normalised to median of control group in each experiment) and frequency of CD69^+^ PD-1^+^ cells within CD44^+^CD8^+^ T cells from tumours, and total CD8^+^ T cells from the blood. **d,** representative gating of IFNγ, TNFα, and IL-2 on respective populations (as in Fig. 5e). Gates based on unstimulated control. **e**, % expression of IFNγ, TNFα, and IL-2 on respective endogenous CD8^+^ (top) and OT-I (bottom) T cell populations. **Statistics: a-e,** pooled from 2 independent experiments, minimum N=5 mice/group/experiment. **b**, one-way repeated-measures ANOVA with Holm-Sidak test, connected points from individual mice. **c**, Two-way ANOVA with Tukey’s test. **e**, one-way repeated-measures ANOVA with Tukey post-test. n.s., p > 0.05; *p<0.05, **p < 0.01; ***p < 0.001; ****p < 0.0001.

**Extended Data Figure 8:**
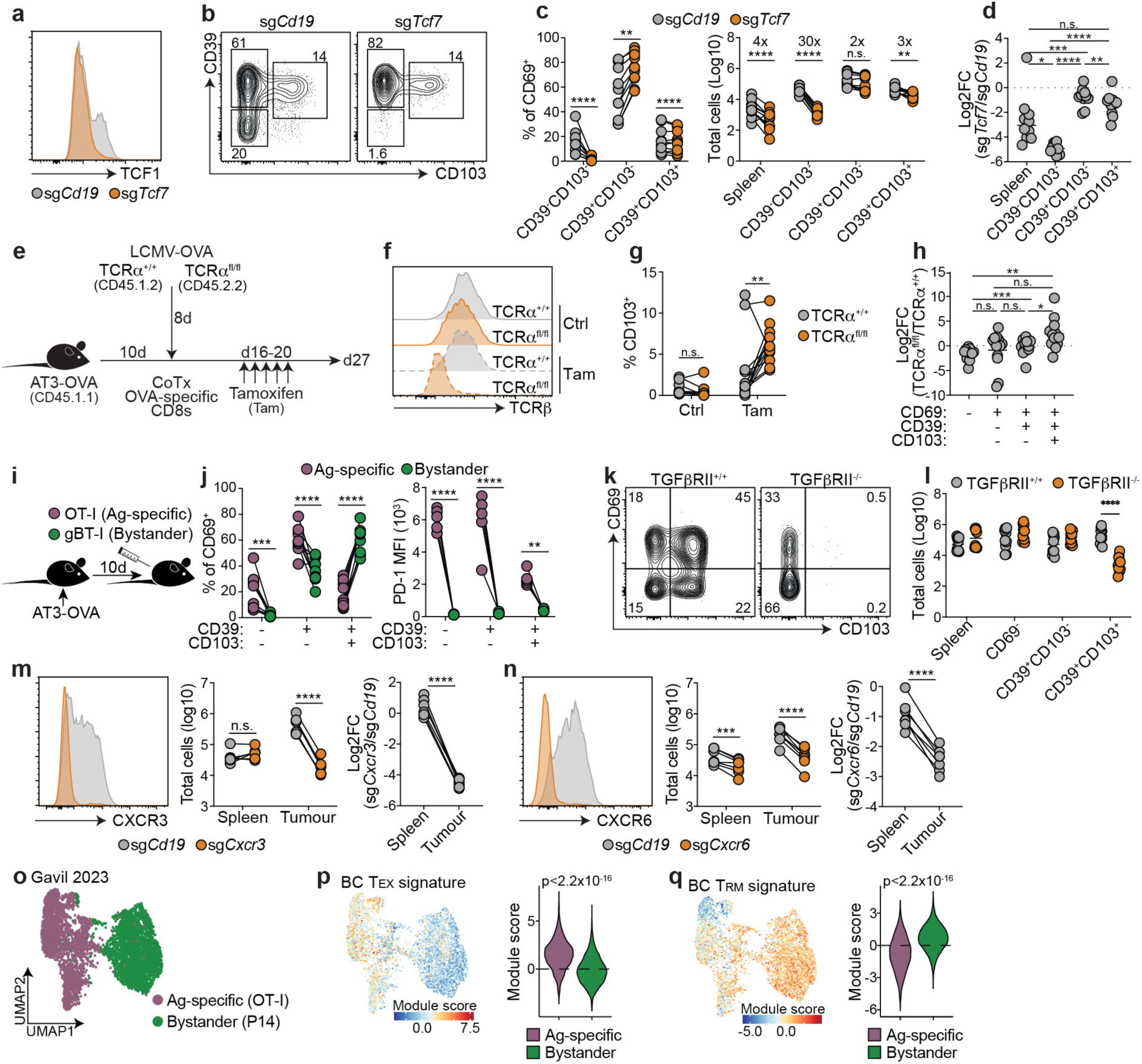
Mechanistic assessment of factors controlling formation of TIL populations. **a-d**, CRISPR/Cas9 depletion of *Tcf7* from naïve OT-I T cells on the development of T cell populations in AT3-OVA tumours harvested 24 d post-inoculation. **a**, TCF1 expression on co-transferred sg*CD19* (control) and sg*Tcf7* treated OT-I T cells isolated from tumours. **b**, Representative gating of CD69^+^ OT-I T cells. **c**, frequency and number of CD69^+^ OT-I T cells from respective gates as in (b). **d**, Log2 fold-change of cell numbers of respective populations. **e-f**, effect of TCRα depletion on TIL populations. Congenically distinct TCRα^+/+^ Cre^ERT2^ and TCRα^fl/fl^ Cre^ERT2^ mice were infected with LCMV-OVA, SIINFEKL-tetramer^+^ CD8^+^ T cells were sorted from spleens at d8 post-infection and co-transferred into AT3-OVA bearing mice 10 d post-tumour inoculation. Mice were then treated with Tamoxifen (Tam) or vehicle control on d16-20 post-tumour, and tumour-derived cells isolated on d27. **e**, schematic. **f**, TCRβ staining on respective populations. **g**, % CD103 expression of respective populations. **h**, ratio of cell counts for respective populations. **i-j**, Effector OT-I (Ag-specific) and gBT-I (bystander) CD8^+^ T cells were co-transferred into mice 10 d post-AT3-OVA inoculation and intratumoural T cells analysed 7 d later. **i**, Schematic. **j**, Frequency and PD-1 expression of respective CD69^+^ populations. **k-l**, Effector control (TGFβRII^+/+^) and TGFβRII^-/-^ bystander OT-I T cells were transferred into AT3 tumour-bearing mice 10 d post inoculation. Representative gating (k) and cell counts (l) of respective populations harvested from tumours 6 d following transfer. **m-n**, Effector control (sg*CD19*) and CXCR3 (m, sg*CXCR3*) or CXCR6 (n, *sgCXCR6*) deficient gBT-I T cells were co-transferred into AT3-OVA tumour bearing mice 10 d post-inoculation and analysed in spleens and tumours 7 d later. Expression of CXCR3 or CXCR6, respective cell counts, and ratios of cells are shown. **o-q,** Analysis of CITEseq of LCMV-specific P14 or tumour-specific OT-I T cells isolated from EO771-OVA BC tumours^16^. **o,** Unified cell-surface protein and RNA-seq expression data represented on weighted nearest neighbours UMAP coloured by T cell transgenic. **p-q,** quantification of BC T_EX_ (**p**) and T_RM_ (**q**) signature module scores, p-value calculated by Wilcoxon signed-rank test. Statistics: **a-d**, i-l, pooled from N=2 independent experiments; m-n, representative of N=2 independent experiments with a minimum of 5 mice per group. **c,g,j,m,n,** linked symbols indicate cells from the same mouse, 2-way repeated-measures ANOVA with Sidak post-test. **d,h,** 1-way ANOVA repeated-measures ANOVA with Tukey post-test. l, 2-way ANOVA with Sidak post-test. p-q, Wilcoxon signed-rank test. p>0.05, *p<0.05, **p<0.01, ***p<0.001, ****p<0.0001.

**Extended Data Figure 9:**
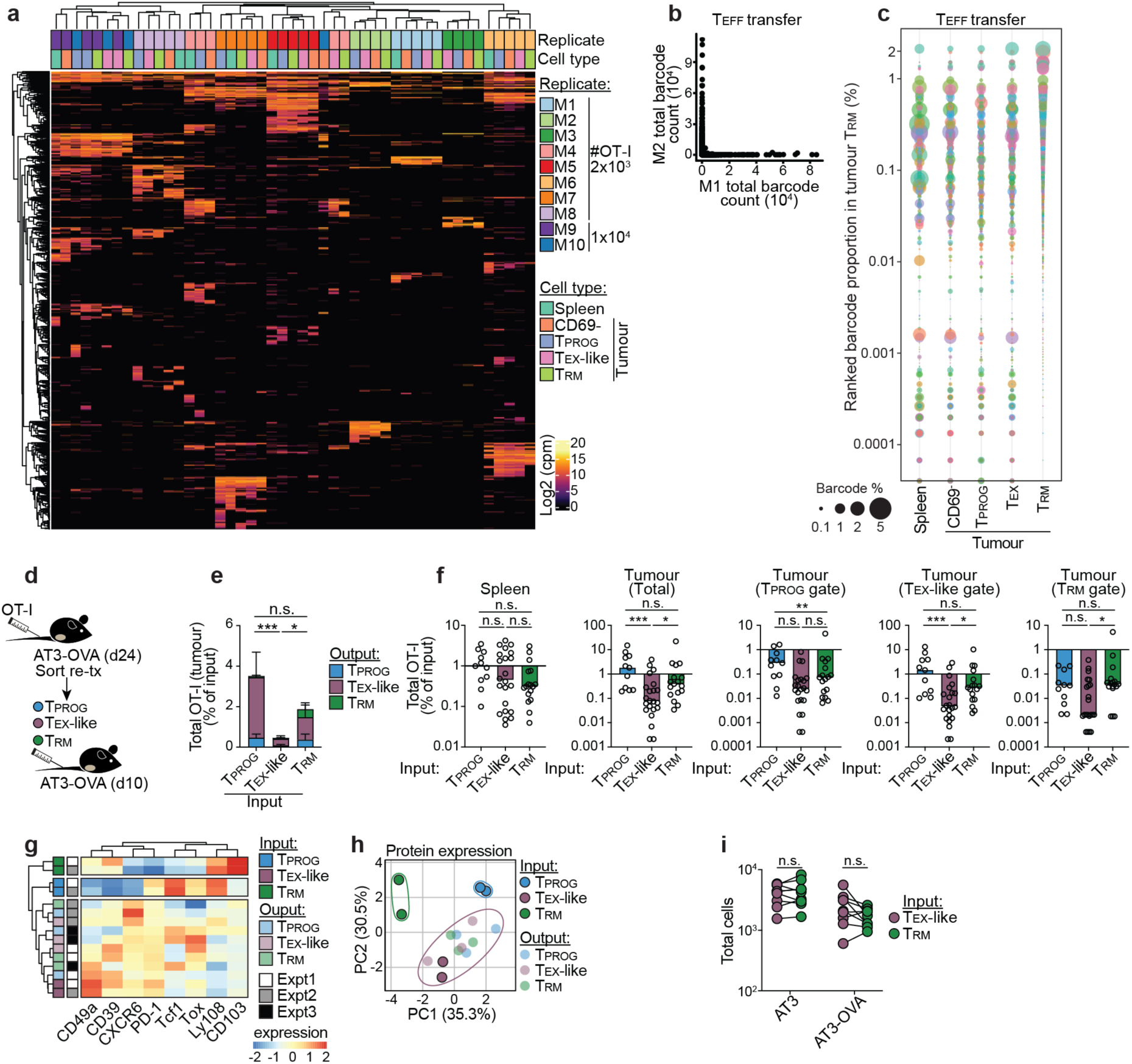
Developmental relationship between intratumoural T_RM_ and T_EX_ cells. **a-c,** Related to Fig. 6a-d. **a**, heatmap showing counts of individual barcodes (y-axis) across different replicate mice and sorted populations from naïve OT-I SPLINTR experiments. **b**, Total barcode-seq counts of each identified barcode from two (M1 vs M2) effector OT-I SPLINTR experimental replicates. **c,** Barcode-seq data from representative effector OT-I SPLINTR experiment (M1) sorted by frequency in the tumour T_RM_ sample, where bubble size reflects clone proportion in sample. **d-h**, Sort and intravenous retransfer of AT3-OVA intratumoural populations. **d,** Schematic. **e,** Enumeration of transferred populations 28 d post-tumour inoculation as the frequency of input cell number. **f**, Cell numbers of respective populations isolated from spleen and tumours, normalised to input frequency, isolated 28 d post-tumour inoculation. **g,** Relative expression of proteins on respective populations of primary transferred (Input) and retransferred (Output) cells 28 d post-tumour inoculation determined by flow cytometry with unbiased hierarchical clustering indicated and **h,** PCA indicating expression of proteins in (**g**), each dot being concatenated samples from independent experiments. **i**, related to Fig. 6e-i, showing total cell counts isolated from tumours at d 22 post-tumour inoculation. Statistics: **a,** pooled from 3 independent experiments, with 10 total replicates (i.e. M1-10). **b,** M1 and M2 indicate independent experiments, each with technical replicates (not shown). **d-h,** Data pooled from 3 independent experiments, N=11-23 mice per group. n.s. p>0.05, *p<0.05, ***p<0.001. **i**, Data representative of 2-independent experiments, linked symbols indicate cells isolated from the same mouse, N=9 mice per group, analysed by 2-way repeated-measures ANOVA with Sidak post-test.

## Methods

### Mice

C57BL/6, gBT-I:CD45.1.2, gBT-I:CD45.1.1, gBT-I:CD90.1.2, P14:CD45.1.2, OT-I:CD45.1.2, OT-I:CD90.1.2; OT-3:CD45.1.2:TCRα^-/-56^, OT-I:TGFβRII^fl/fl^:Cre^dLck^:CD45.1(TGFβRII^-/-^) and TCRα^fl/fl^Cre^ERT2^ female mice were bred and maintained in the Department of Microbiology and Immunology, University of Melbourne under a 12h/12h light/dark cycle, at 19-22°C and 40-70% humidity. All experiments were approved by the University of Melbourne Animal Ethics Committee (ID nos. 21651 and 21938). All mice were between 6-14 weeks of age at the beginning of the experiments.

### Human studies

This project was approved by the Human Research Ethics Committee of the University of Melbourne (ID nos. 13009 and 14517). All participating patients provided written informed consent.

### Isolation of lymphocytes from human tumours and tissues

Following tumour excision, a representative tumour fragment was processed to generate single-cell suspensions as previously described^1^. Briefly, tumour or healthy tissues were finely diced in RPMI1640 containing 10% FCS and 0.5mg/ml collagenase D (Worthington Biochemical, Lakewood, NJ) and were incubated for 30 min at 37°C. Digested pieces were mashed through 70-μm strainers and washed with RPMI1640 with 10% FCS. Lymphocytes from liver tumours and healthy liver tissues were enriched through density gradient centrifugation (500*g*, 20min at 25°C) on a 44% / 70% isotonic Percoll gradient (GE Healthcare, diluted in HBSS, and cells at the solution interphase were isolated and washed with HBSS. Blood was diluted 1:1 in HBSS, overlaid on Ficoll-Paque PLUS (Sigma Aldrich), centrifuged (400g, 15min, 25°C), and peripheral blood mononuclear cells (PBMCs) isolated from interphase. Cells were either utilised immediately for flow cytometry, or frozen in 10% DMSO: 90% FCS freezing media (breast, breast tumour, and blood samples) or Cryostor CS10 Freeze Media (StemCell Technologies, #07930; liver and liver tumour samples) for CITEseq and restimulation assays.

### Murine tumour and infection models

The murine TNBC cell line, AT3-OVA was provided by Prof. Phillip Darcy (Peter MacCallum Cancer Centre, Melbourne, VIC, Australia)^57,58^. AT3-OVA cells were cultured with complete DMEM (DMEM, 10% FCS, 2 mM L-glutamine, 100U ml^-1^ penicillin, 100mg ml^-1^ streptomycin). 5×10^5^ AT3-OVA cells in exponential growth phase (∼70-80% confluency) were injected orthotopically into the 4^th^ mammary fat pad in a total volume of 50µl HBSS.

The murine melanoma B16F1-gB.GFP (B16-gB) cell line was provided by Jason Waithman (University of Western Australia)^47^. B16-gB cells were cultured and passaged in complete RPMI (RPMI1640, 10% FCS, 2mM L-glutamine, 100U ml^-1^ penicillin, 100mg ml^-1^ streptomycin, 50mM 2-mercaptoethanol) at 37°C and 5% CO_2_. B16-gB was inoculated epicutaneously as previously described^47^. Tumours were measured every 2-3 days following the development of palpable tumours using vernier callipers, and tumour volume was calculated (length × width^2^)/2. Mice were euthanised when tumours reached an ethical limit of 1.0 cm^3^. Effector gBT-I or OT-I T cells were transferred into mice following the observation of tumour growth, between 14-20 d following tumour inoculation, and then harvested at tumour endpoint (when tumours reached a maximum of 1000mm^3^) which occurred 14-20 d following T cell transfer.

LCMV infection was performed by intraperitoneal (i.p.) injection of 2×10^5^ pfu of the Armstrong strain of LCMV.

### T cell transfer

For the adoptive transfer of naïve T cells, 1-5×10^4^ transgenic (P14 or OT-I) T cells were transferred intravenously to recipient mice 1-2 d before infection with LCMV or inoculation with AT3-OVA. For effector T cell transfer, transgenic (P14, OT-I, OT-3, or gBT-I) T cells were activated in culture for 5 d with gp_33-41_ (KAVYNFATM – P14; Auspep), OVA_257-264_ (SIINFEKL – OT-I/OT-3; Auspep), or gB_498-505_ (SSIEFARL – gBT-I; Auspep) peptide-pulsed splenocytes, in the presence of recombinant human IL-2 (25 U ml^-1^; Peprotech) in complete RPMI (as above) at 37°C and 5% CO_2_. T cells were split 1:1 with fresh media and IL-2 on days 2-4. T cells were resuspended in 200µl HBSS for intravenous transfers. Unless stated otherwise, 1×10^4^ effector OT-I, 5×10^6^ gBT-I, or 1×10^6^ OT-3 were transferred 10 d post-AT3-OVA inoculation.

### *In vivo* treatments

For *in vivo* intravascular staining, 3μg of αCD90.2-biotin was injected i.v. in 200μL PBS, 3min before euthanasia. To inhibit S1P-signaling pathways, mice were administered FTY720 (Cayman Chemical) diluted in 2% (2-Hydroxypropyl)-β-cyclodextrin (Sigma-Aldrich) or vehicle daily via i.p. injection (1μg/g) for the indicated times in the figure legend (Extended Data Fig. 7c). For tamoxifen treatment, mice were administered 2 mg of tamoxifen (or ethanol as vehicle control) diluted in sunflower seed oil (both from Sigma) i.p. daily for a total of five injections.

### Isolation of lymphocytes from mouse tissues

Lymphocytes from spleens and lymph nodes were isolated by grinding through 70-μm strainers. AT3-OVA tumours and MFPs (avoiding the inguinal lymph node) were collected into collagenase III solution (3mg ml^-1^; Worthington) containing DNase I (2.5 mg ml^-1^; Sigma), chopped into fine pieces and incubated for 60 min at 37°C. B16-gB tumours were collected into liberase TL research grade solution (0.25 mg ml^-1^; Roche), chopped into fine pieces, and incubated for 60 min at 37°C. Spleens and tumours were passed through a 70-mm strainer and erythrocytes were lysed in red cell lysis buffer (eBioscience) prior to staining for flow cytometry. Cells isolated from B16-gB tumours were cryopreserved in 10% DMSO:90% FCS for scRNAseq.

### Flow cytometry and cell sorting

Single-cell suspensions were stained with fluorescently conjugated antibodies at 4°C for 30-45 min in FACS buffer (1% BSA and 0.05M EDTA in PBS). Dead cells were excluded by staining with Zombie Aqua or Near Infrared dyes (Biolegend). For transcription factor and cytokine staining, samples were fixed and permeabilised using the Foxp3/transcription-factor-staining buffer kit (eBioscience) according to the manufacturer’s instructions and stained with fluorescent antibodies against intracellular proteins in permeabilisation buffer (containing 2% rat and mouse serum (eBioscience)). Cells were enumerated using SPHERO calibration particles (BD Biosciences). Fluorescently labelled cells were acquired on a Cytek Aurora, unmixed with SpectroFlo® software and analysis was performed using FlowJo (v.10.10.0; Treestar), or OMIQ for high-dimensional flow cytometry analysis. For cell sorting experiments, cells were sorted using a BD FACS Aria, using a 100 μm nozzle. For human cell sorting experiments for CITEseq, CD3^+^ Zombie NIR^-^, cells were sorted into 50% FCS in RPMI before downstream processing. For B16-gB sorting experiments, DAPI^-^ OT-I or gBT-I cells were sorted into 50% FCS in RPMI before downstream processing. For mouse AT3-OVA sort-transfer experiments, mice were treated with anti-ARTC2 nanobody intravenously (50 mg per mouse; S+16a, Biolegend) 10 minutes before organ collection, stained as above, and respective populations (see figure legend) sorted into 50% FCS in RPMI before washing with HBSS and transfer into recipient mice (8×10^4^ cells in 200 µl intravenously).

### *In vitro* stimulation assays

To assess cytokine production capacity by T cells, cells were stained with surface stain antibodies (as above), then incubated with phorbol myristate acetate (PMA; 50 ng ml^-1^; Sigma-Aldrich), Ionomycin (1 mg ml^-1^), Brefeldin A (10 mg ml^-1^, Sigma-Aldrich), GolgiStop (1:1500, BD) in complete RPMI (as above) for 4-5 h before intracellular staining and flow cytometry. Unstimulated controls were included to confirm that stimulation did not alter cell surface staining and to serve as negative controls for staining.

### Bulk RNA-seq analysis

(For Fig. 1a) Scatter plot of logFCs (log2-fold-changes) was produced using raw RNA-seq count data from GEO for Man et al. (accession GSE84820^26^) and Mackay et al. (accession GSE70813^8^). From the Man data, wild-type day 30 chronic and acute LCMV samples were selected for analysis, while from the Mackay data, the following samples were selected for three separate analyses: (1) Skin T_RM_, T_CM_, and T_EM_ HSV samples; (2) Gut T_RM_, T_CM_, and T_EM_ LCMV samples; and (3) Liver T_RM_, T_CM_, and T_EM_ LCMV samples. T_CM_ and T_EM_ samples were subsequently treated as a single ’T_CIRC_’ group. For each analysis, genes were annotated with information from NCBI and those with obsolete symbols or annotated as rRNA were removed, as were genes that failed to achieve a count above 10 in all samples in at least one group. Each dataset was further processed by applying the imputation strategy published previously^59,60^. Counts-per-million values were calculated using the *edgeR* package^61^, together with scaling factors derived from the TMM method^62^, log2 transformed with a prior count of 1, followed by application of the normalisation method RUV-III^63^ with biological replicates nominated as replicates, mouse housekeeping genes^56^ nominated as ‘negative control’ genes, and k=1 factors of unwanted variation. Normalisation success was assessed with relative log expression plots^64^, PCA plots^65^, and histograms of p-values. The *limma* package^66^ was used to fit gene-wise linear models for the given group structure with the output from RUV-III as an additional model covariate. LogFC estimates from the Mackay Skin, Gut, and Liver analyses were averaged to create a ’pooled T_RM_’ expression profile and then plotted against the estimates from the Man analysis. A p-value was calculated by constructing a two-way contingency table, counting the number of genes with concordant/discordant logFCs, and applying Fisher’s exact test for association. Core T_RM_ gene signature was obtained as described previously^67^.

### Single-cell CITE/RNA sequencing library preparation

Following sorting of cell populations, sorted cells were stained with TotalSeq-C Universal Cocktail V1.0 (Biolegend; Human CITEseq experiments) or TotalSeq-C Hashtag antibodies (Biolegend: mouse B16-gB scRNA-seq experiment) according to manufacturer’s instructions. Cells were filtered through a 40-mm Flowmi cell strainer, loaded onto a 10X chromium controller, and prepared for sequencing using a Chromium Next GEM Single Cell 5’ kit with feature barcoding and immune receptor mapping (v2, Dual Index) and VDJ enrichment kits for mouse or human TCR from 10X. Libraries were generated according to the manufacturer’s instruction. Libraries were profiled on an Agilent Tapestation and quantified using a Qubit before sequencing on an Illumina NextSeq 2000 P2 or P3 kit.

### Single-cell CITE/RNA sequencing analysis

Sequencing reads of three separate lanes were aligned to the hg38 reference genome and T cell receptor reference (VDJ) and counted with cell ranger-6.1.2. CITEseq antibodies were aligned to their custom reference sequences from BioLegend. Patient samples were demultiplexed into genotypic donors using vireo^68^ on aligned BAM files from three lanes. Donors were matched to genotypes using known cell frequencies in each sample. Three batches of single-cell data were merged and processed using Seurat^69^ (version 4.3.0). Specifically, cells were filtered if they contained fewer than 500 genes, more than 5% mitochondrial RNA, and were annotated to have more than 1 beta chain in the VDJ assay or two genotypes in the vireo analysis (cell doublets). Then, for the RNA assay, we used NormalizeData to normalise the counts data and determine the top 2000 variable genes using FindVariableFeatures. We excluded T cell receptor components (^TRA/^TRB), mitochondrial genes (^MT), and ^HLA from the variable genes as unwanted factors of variance and performed principal component analysis (RunPCA). The CITEseq assay was processed similarly with CLR normalisation and margin=2. We removed the individual effects between the donors using Harmony^70^ on the RNA and CITEseq assays individually and then combining the correction reductions using the FindMultiModalNeighbors functions (using 25 dimensions from the RNA reduction and 18 from CITEseq). UMAP reductions and cell neighbours were calculated using 25 dimensions from the weighted nearest neighbour reduction. Clusters were detected with a resolution of 3. We then determined T cell subsets within the data by assessing cluster markers (FindAllMarkers) and their overall expression and comparing them to published markers in the literature (merging clusters as need be). We subset the data to CD8+ T cells only at this point by excluding other T cell populations and contaminating immune populations. All further analyses and plots were generated in R (version 4.2) using tidyverse^71^ functions and ShinyCell^72^. Heatmaps were created with the pheatmap package (version 1.0.12). Subset signature expression levels were calculated using the AddModuleScore function. Pseudobulk analysis was performed using the edgeR package^61^: counts of reads were summed per subset. Pseudobulk samples from the periphery of the data (gdT, MAIT) are filtered from the analysis and lowly expressed genes removed using filterByExpr(). The samples were then normalised using TMM and the calcNormFactors() function. Dimensional reduction was performed using the plotMDS function. Other single-cell datasets, such as Giles 2022^27^, or subsets of the data (tumour only) were analysed analogously to the methods above (leaving out CITEseq and TCR where appropriate).

#### TCR analysis

TCRs from cellranger outputs were paired based on cell barcodes and merged with gene expression data. In cases where multiple contigs were detected the contigs with the highest UMI was kept. TCR clonotype was defined by the joined alpha and beta CDR3 nucleotide sequences, and expanded TCR clonotypes were determined by filtering the list of clonotypes to those that are found in 2 or more cells. Antigen specificity of cells was based on TCR clonotype and donor HLA identity. HLA type was determined with *ArcasHLA* using the ‘—single’ flag^73^. Reads covering the genome co-ordinates chr6:28510120-33480577 were extracted into a separate fastq file per donor and processed individually. Plots of clonotype diversity and similarity within/across celltypes were calculated using djvdj (https://rnabioco.github.io/djvdj/ version 0.1.0) and plotted with ComplexHeatmap^74^. For clonotype sharing plots, expanded clonotypes within each tissue were filtered to only include those which are shared across more than one subset. The number of shared clonotypes was quantified for each subset pairing, and the total number of cells contributing to clonotype sharing from each subset was counted. Visualisations were generated with custom R scripts using the *ggraph* package (version 2.0.5). For iterations of the sharing plots with filtering, input data were refined so that a shared TCR was removed from the plot if one of the celltypes in the pairing contributed just 1 cell (for n>1 filtering) or 2 or fewer cells (for n>2 filtering). Other visualizations of clonotype cell fate and cluster sharing were generated using custom R scripts, as provided. Comparisons between TCR sequences was performed using TCRdist3^33,34^ using default weights with the distance matrix calculated as the sum pairwise distance for the alpha and beta chains. To determine potential virus-recognising clones, published TCRs with known viral HLA-peptide specificities were obtained from VDJdb^35–37^ following which edit distances were calculated for each clone against reference CDR3a/b peptide sequences. The lowest edit distance was obtained for each clone, with distances less than one considered viral-associated.

#### Pan-cancer atlas processing^28^

Preprocessed Seurat objects were obtained^28^. SCTransform was used to normalise, scale and regress pre-filtered dataset. Pre-computed scores for dissociation induced genes(DIG), Malat1, mitochondrial percentage and cell-cycle(G2M and S scores) were used for regression. T-cell receptor variable/joining (^TR[A|B|G|D][V|J]), B-cell receptor variable/joining (^IG[H|L|K][V|J|D]) DIG(^HSP/^DNAR), cell-cycle, DIG(^HSP/^DNAJ) and ribosomal (^RP([0-9]+-|[LS])) genes were removed from VariableFeatures gene list priors to calculating principal components(PC). Subsequent UMAP and FindNeighbours commands performed with 15 PCs. T_RM_ and T_EX_ scores were calculated with the AddModuleScore command using respective genesets. T_EM_ and T_EMRA_ scores were calculated using differentially expressed genes calculated from the BC dataset.

### Signature acquisition and Module Scoring

T_RM_ and T_EX_ expression signatures were derived by summing raw counts, gene-wise, over each donor within each cluster, to create ‘pseudo-bulk’ samples for each donor/cluster combination. Samples composed of less than 20 cells were removed, and samples from the same meta-cluster were subsequently treated as belonging to the same group. Genes annotated as being of ’HLA’, ’TRB’, ’TRA’, ’TRG’, or ’TRD’ type were removed, as were genes that failed to achieve a count above 5 in at least 2 samples in at least one group (breast) or a count above 3 in at least 4 samples in at least one group (liver). Counts were log2 transformed with a prior count of 1/2 (breast) or 1 (liver), followed by application of the normalisation method RUV-III with samples from the same group nominated as replicates, human housekeeping genes nominated as ‘negative control’ genes, and k=20 (breast) or k=15 (liver) factors of unwanted variation. Normalisation success was assessed as above. The edgeR package [*] was used fit gene-wise negative binomial generalized linear models for the given group structure with the output from RUV-III as additional model covariates, with a prior count of 1/2 (breast) or 1 (liver). Likelihood ratio tests were employed to test for differential expression (DE), where a gene was judged to be DE if the Benjamini and Hochberg^75^ adjusted p-value < 0.05. For breast, the T_RM_ signature was defined by a contrast between the T_RM_ group and the average of all other groups; the T_EX_ signature was defined similarly. For liver, the T_RM_ signature was defined by a contrast between the CD103^+^ T_RM_ group and the average of all other groups except the CD103^-^ T_RM_ group; the T_EX_ signature was defined similarly. The breast T_RM_ ‘union’ signature was derived using the same steps for breast above, except that samples obtained from the T_RM_ or T_EX_ clusters were combined into a single group. Signature scores were calculated from the SCTransformed assay using the ’AddModuleScore’ function from the Seurat package^69^ for up and down genes separately and combined by averaging the scores for the up genes and the sign reversed scores for the down genes. These scores were overlayed onto UMAP plots using the FeaturePlot function from Seurat. To generate the pan-cancer T_RM_ signature we focused on the leading-edge genes contributing to the enrichment in the barcode plots (Fig. 3c, Extended Data Fig. 4g). Specifically, the leading-edge genes contributing to T_RM_ signature associations between the BC and liver datasets were refined by intersecting the genes in the breast and liver T_RM_ signatures (described above), then subsetting to genes which are either (1) concordantly DE in both breast and liver or (2) DE in one with a concordant absolute logFC > 0.5 in the other. The pan-cancer T_EX_ signature was refined similarly (Extended Data Fig. 4g). Barcode enrichment plots were generated using *limma*, and gene set enrichment p-values were calculated using the camera function on the log fold-changes for populations of interest against the background calculated with FindMarkers.

### Survival Analysis

(For Fig. 2l, Extended Data Fig. 3f-h) Clinical information and normalised microarray data for the study^50,76^ were downloaded from the cBioPortal (https://www.cbioportal.org). Signature scores for each patient were calculated using the ‘sig.score’ function from the *genefu* package^77^. Kaplan-Meier curves were calculated using the R package *survival* and were plotted, with the log-rank test p-value, using the R package *survminer*.

(For Fig. 2m) Clinical information and microarray data for the iSPY study^31^ were downloaded from supplementary material and GEO (accession GSE194040), respectively. Probes targeting the same gene were averaged. If a gene had missing values, these were imputed using the average of all non-missing values across the gene. For each platform batch, relative log expression^64^ values were computed, the median value for each sample was calculated, then samples were ranked by this median value: the bottom, middle, and top third rankings defined 3 separate ‘pseudo-batches’ of samples. For each PAM50 breast cancer expression subtype in each pseudo-batch, one of the following was performed: if there were between 5-10 samples, 5 samples were randomly selected and averaged, gene-wise, to create one ‘pseudo-sample’; or if there were >10 samples, the bottom 5 ranked samples and the top 5 ranked samples were averaged, gene-wise, the create 2 pseudo-samples. These pseudo-samples were used as pseudo-replicates in the RUV-III-PRPS normalisation methodology^78^, with all genes nominated as ‘negative control’ genes, and k=7 factors of unwanted variation. Normalisation success was assessed with relative log expression and PCA plots^65^. Based on this normalised data, signature scores for each patient were calculated using the ‘sig.score’ function from the *genefu* package. ROC curves and p-values were calculated using the R packages *pROC*^79^ and *verification*, respectively.

### Cyclic immunofluorescence (CycIF) of human breast tumours

#### Selection of breast cancer samples

FFPE-embedded tumour tissues from 7 patients (6 female, 1 male) were purchased from the Cooperative Human Tissue Network (CHTN) based on histologic criteria, for invasive carcinoma (ductal/lobular) and 5 µm thick sections were cut on Superfrost Plus histology slides (Fisher scientific) at the BWH Histopathology core, as previously described^29^.

#### CycIF staining

FFPE tissues were deparaffinized, rehydrated, subjected to antigen retrieval on a Leica BondRx and characterised by cyclic immunofluorescence (CycIF) imaging, as previously described^29^. Briefly, tissues were stained overnight at 4°C with primary antibodies or for 1 hour at RT for secondary antibody conjugates in a dark, humidified chamber with antibodies diluted in Superblock (Thermo-Fisher) supplemented with 1 mg/ml Hoechst 33258 (Bio-Rad Laboratory) rinsed for 30 minutes in PBS (RT) and mounted with 50% glycerol in PBS using a 24×60mm coverslip. All samples were stained and imaged in together to reduce batch effects using the antibody panel. Stained tissues were imaged using a Cytefinder II HT (Rarecyte) automated slide scanning fluorescence microscope using a 20x (0.6 NA) objective. After imaging, mounted slides were soaked in PBS (RT) to detach coverslips, then immersed in PBS supplemented with 4.5 % hydrogen peroxide and 24 mM sodium hydroxide and exposed to LED light for 1 hour. Slides were then rinsed twice in PBS in preparation for the next staining cycle. Image processing, assembly, segmentation and single-cell quantification were performed using MCMICRO^30^.

#### CycIF image analysis

CycIF Images were processed with Cecelia^80^. Individual channels were denoised and segmented with Cellpose^81^ using a radius of 10 and 8 μm, respectively. T cell (CD3e) and cancer cell (panCK) segmentation was based on their respective markers in combination with the DNA (Hoechst-stained nuclei) channel from the third imaging cycle, the same cycle as CD3e. Cell populations were gated using Cecelia’s histocytometry module and spatial interactions analysed. Cell distances were extracted using K-nearest neighbour from the DBSCAN library^82^. Clusters within the CD8^+^CD103^+^KLF2^-^ T cells were defined using the Seurat package. We first removed the individual donor effects and performed principal component analysis using a set of 22 features (CD39, CD45RO, CD3E, CD25, CD73, CD8A, GZMB, CD103, LAG3, PD1, TIM3, CD69, TCF1, FOXP3, GNLY, CD4, pJUN, pERK, KLF2, CD7, CD94, NKG2A) (RunPCA). Shared neighbour graphs (FindNeighbors) and UMAP reduction (RunUMAP) were performed using the first 10 dimensions. Clusters were detected using a resolution of 0.8 (FindClusters).

### CRISPR/Cas9 editing of CD8^+^ T cells

CRISPR/Cas9 editing of naïve CD8^+^ T cells was performed as previously described^83^. Single guide RNAs (sgRNA) targeting: *Tcf7* (5′-UCUGCUCAUGCCCUACCCAC-3′, 5′-AGCUGGGGGACGCCAUGUGG-3′, 5′-UGUGCACUCAGCAAUGACCU-3′), *Cxcr3* (5’-GAACAUCGGCUACAGCCAGG-3’, 5’-UGAGGGCUACACGUACCCGG-3’), *Cxcr6* (5’-UCUGUACGAUGGGCACUACG-3’, 5’-UGUGCCAAAGACCCACUCAU-3’), and *Cd19* (5′-AAUGUCUCAGACCAUAUGGG-3′, 5’-GAGAAGCUGGCUUGGUAUCG-3’) were purchased from Synthego (CRISPRevolution sgRNA EZ Kit). sgRNA/Cas9 RNPs were formed by incubating 0.3 nmol of each sgRNA with 0.6 μl Alt-R S.p. Cas9 nuclease V3 (10 mg ml^−1^; Integrated DNA Technologies) for 10 min at room temperature. Naïve CD8^+^ T cells were negatively enriched from spleen and lymph nodes of gBT-I or OT-I mice by incubating cell suspensions with anti-CD4, anti-CD11b anti-F4/80, anti-Ter119 and anti-I-A/I-E monoclonal antibodies, followed by incubation with goat anti-rat IgG-coupled magnetic beads (Qiagen) before removing bead-bound cells. 1×10^7^ enriched T cells were resuspended in 20 μl of P3 (P3 Primary Cell 4D-Nucleofector X Kit; Lonza), mixed with sgRNA/Cas9 RNP and electroporated using a Lonza 4D-Nucleofector system (DN100). Cells were rested for 30 min in a 96-well plate before direct transfer into recipient mice (co-transfer of 1:1 ratio of sg*Cd19*:sg*Tcf7* CRISPR-edited naïve gBT-I cells, total of 2 × 10^4^ cells per mouse: Extended Data Fig. 8a-d) or activation in culture for 5 days with peptide pulsed splenocytes (gB_498-505_ (SSIEFARL)) in the presence of IL-2 (25 U ml^−1^, Peprotech) at 37 °C, 5% CO_2_, before transfer into recipient mice (co-transfer of 1:1 ratio of sg*Cd19*:sg*Cxcr3 or* sg*Cd19*:sg*Cxcr6* CRISPR-edited naïve gBT-I cells, total of 1×10^6^ cells per mouse: Extended Data Fig. 8m-n).

### SPLINTR Methods

#### Production of libraries

To construct the SPLINTR barcoding systems, Violet-light excited GFP (VEX) was used to replace NGFR in MSCV-IRES-NGFR^84^ using the NcoI and BamH1 restriction sites, or eGFP into the SPLINTR lentivirus vector^41^. A semi-random oligonucleotide library synthesised by Integrated DNA Technologies with the following pattern (NNSWSNNWSW)_6_ was amplified by eight cycles of PCR. The barcode library was cloned into the 3’UTR of VEX using BamH1 and MfeI restriction sites at a 50:1 insert:vector ratio. 10 side-by-side ligation reactions were pooled and purified using two Monarch PCR & DNA Clean up columns (NEB) in a volume of 6 µl per column. The ligation reactions were pooled and split across two 25 µl aliquots of Endura Electrocompetent cells (Lucigen). Cells were recovered for 1 hr post-electroporation, pooled and grown in 500 mL of LB supplemented with ampicillin (100 µg ml^-1^) overnight at 37°C. The plasmid library was extracted using NucleoBond Xtra kit (Macherey-Nagel). SPLINTR libraries were sequenced to a depth of 100 million paired-end reads per technical duplicate and reference libraries used for downstream analysis were generated as described previously^41^. SPLINTR retrovirus and lentivirus VEX library represents a highly diverse barcode library containing 3.2×10^6^ unique barcodes or 1.3×10^6^ barcodes, respectively.

#### Generation of SPLINTR^+^ OT-I T cells

Naïve (T_N_) OT-I T cells were barcoded via transduction of hematopoietic stem and progenitor cells (HSPC) and intra-thymic transfer into sublethally irradiated chimeric mice. Briefly, bone marrow was isolated from OT-I mice, and lineage(lin)-positive cells were depleted using the EasySep™ Mouse Hematopoietic Progenitor Cell Isolation Kit (Stem Cell Technologies). Lin-negative HSPCs were plated in fibronectin-coated 12-well plates containing polyvinyl alcohol (PVA)-based media: 1× Ham’s F-12 Nutrient Mix liquid media (Gibco) supplemented with 10 mM HEPES, 1× P/S/G, 1 ITSX, 1 mg/mL PVA along with TPO (100 ng/µL) and mouse SCF (10 ng/µL). For barcoding, 20 pools of 1×10^6^ cells (cultured for 6 d) were transferred to 24-well fibronectin-coated plates in 250 μL of PVA media. Viral barcoding vectors were titrated to achieve 10% reporter expression (0.1 MOI) to minimize multiple integrations. Cells were transduced via spinfection at 2000g for 2 h (no break). After transduction, media was refreshed and cells were cultured for a further 48 h before VEX-positive and Lin-cells were sorted (ARIA Fusion, BD). The 20 pools of 3.5-5.5×10^5^ cells were individually plated into 48-well plates and expanded for 1-week. Recipient C57Bl/6 mice were then irradiated 4Gy and HSPCs were transplanted intrathymically such that each mouse received an independent barcode pool. 8 weeks later, chimeric mice were bled, equal numbers of VEX^+^ OT-I T cells from each mouse were pooled, and cells were transferred into C57Bl/6 mice that were then inoculated with AT3-OVA tumours.

Effector (T_EFF_) OT-I T cells were barcoded as follows. Naïve OT-I T cells were isolated from spleens and LNs and enriched via negative selection as for CRISPR/Cas9 experiments. Enriched OT-I T cells were activated with anti-CD3 (2μg ml^-1^, 145-2C11, BioXCell) and anti-CD28 (1μg ml^-1^, 37.51, BioXCell) for 24 hours before ‘spinfection’ with a pre-titrated volume of SPLINTR retrovirus to ensure <5% (0.05 MOI) transduction efficiency to limit multiple barcode integrations in a single cell, in 24-well plates, pre-coated with Retronectin (32 ug ml^-1^, Takara). Transduced cells were sorted 24 hours later and transferred into mice bearing AT3-OVA tumours. At the endpoint, transduced OT-I T cell populations were sorted from spleens and tumours and lysed in Viagen lysis buffer with 0.5 μg/ml proteinase K (Invitrogen) for DNA barcode sequencing.

#### Sequencing and analysis

Barcode sequences were amplified from genomic DNA using primers flanking the constant region of the barcode before adding i5 and i7 indexes compatible with NGS sequencing^41^. Libraries were sequenced on an Illumina NextSeq2000 using 100 bp single end chemistry targeting 2 million reads per sample. The *BARtab* pipeline (https://github.com/DaneVass/BARtab)^85^ was used to map the sequencing reads to a barcode reference library, perform qc analysis, and generate a barcode counts table. The *bartools* R package^85^ was used to collapse PCR replicates and generate barcode abundance bubble plots and correlation heatmaps for data visualisation. Scatterplots comparing barcode abundance between two samples were generated using the R package *ggplot2*.

## Data and material availability statement

All data are available from the corresponding author upon reasonable request. The single-cell CITE and RNA-sequencing data generated from this study will be deposited in the GEO before publication. Source data are provided with this paper.

## Code availability

The code generated and used for the analysis of single-cell CITE and RNAseq data is provided with this paper: https://github.com/CSI-Doherty/Burn2025.

## Ethics approval

All animal experiments were approved by The University of Melbourne Animal Ethics Committee (ID nos. 21651 and 21938). All study on human specimens was approved by the Human Research Ethics Committee of the University of Melbourne (ID nos. 13009 and 14517). All participating patients provided written informed consent.

## Acknowledgements

We thank the Flow Cytometry Unit and Bioresources Facility at the Doherty Institute (University of Melbourne) for technical assistance. We thank all Pfizer Inc. Cancer Immunology team members, the laboratory of L.K.M, Professor William Heath, and Dr. Yannick Alexandre (University of Melbourne) for comments and discussion. This work was supported by Pfizer Inc., Cancer Council Victoria Grants-in-Aid, and National Health and Medical Research Council (NHMRC) to L.K.M. L.K.M. is a Senior Medical Research Fellow supported by the Sylvia and Charles Viertel Charitable Foundation. Z.M. is supported by NCI grant R50-CA252138. Histopathology was supported by NIH core grant P30-CA06516.

## Ethics declarations

The authors declare no competing interests.

## Notes

### Competing Interest Statement

The authors have declared no competing interest.

